# Recruitment of the SNX17-Retriever recycling pathway regulates synaptic function and plasticity

**DOI:** 10.1101/2023.02.20.529299

**Authors:** Pilar Rivero-Ríos, Takao Tsukahara, Tunahan Uygun, Alex Chen, Garrett D. Chavis, Sai Srinivas Panapakkam Giridharan, Shigeki Iwase, Michael A. Sutton, Lois S. Weisman

## Abstract

Trafficking of cell-surface proteins from endosomes to the plasma membrane is a key mechanism to regulate synaptic function. In non-neuronal cells, proteins recycle to the plasma membrane either via the SNX27-Retromer-WASH pathway, or via the recently discovered SNX17-Retriever-CCC-WASH pathway. While SNX27 is responsible for the recycling of key neuronal receptors, the roles of SNX17 in neurons are less understood. Here, using cultured hippocampal neurons, we demonstrate that the SNX17 pathway regulates synaptic function and plasticity. Disruption of this pathway results in a loss of excitatory synapses and prevents structural plasticity during chemical long-term potentiation (cLTP). cLTP drives SNX17 recruitment to synapses, where its roles are in part mediated by regulating surface expression of β1-integrin. SNX17 recruitment relies on NMDAR activation, CamKII signaling, and requires binding to the Retriever and PI(3)P. Together, these findings provide molecular insights into the regulation of SNX17 at synapses, and define key roles for SNX17 in synaptic maintenance and in regulating enduring forms of synaptic plasticity.

## Introduction

Cell surface proteins regulate critical cellular functions, including cell adhesion, nutrient uptake and signal transduction. These proteins are regulated in part by their controlled removal from the cell surface via endocytosis, from there, they are either targeted for recycling back to the plasma membrane or routed to lysosomes for degradation (Cullen and Steinberg, 2018; Naslavsky and Caplan, 2018). In neurons, the surface expression of neurotransmitter receptors, ion channels, and adhesion molecules critical for synaptic function depends on precise endomembrane trafficking mechanisms (Kennedy and Ehlers, 2006; Hiester et al., 2018) and defects in the regulated trafficking of these proteins contribute to neurological disease.

To date, two major recycling pathways from endosomes to the plasma membrane have been discovered. One relies on the sorting nexin SNX27 and the well-established Retromer complex. SNX27 is the cargo adaptor, and binds to multiple transmembrane proteins through its N-terminal PSD-95/Dlg/ZO-1 (PDZ) domain (Steinberg et al., 2013). SNX27 cargoes include key neuronal receptors, such as AMPA-type glutamate receptors (Loo et al., 2014; McMillan et al., 2021), β-adrenergic receptors (Lauffer et al., 2010; Temkin et al., 2011), Kir3 potassium channels (Lunn et al., 2007), the 5-hydroxytryptamine 4a (5-HT4a) receptor (Joubert et al., 2004) and the N-methyl-D-aspartate receptor 2C (Cai et al., 2011). In addition to its cargo binding role, the SNX27 PDZ domain binds to the Retromer subunit VPS26 (Gallon et al., 2014). The Retromer complex is composed of three subunits: VPS35, VPS26 and VPS29, which form a 1:1:1 heterotrimer (Hierro et al., 2007). VPS35 forms a central platform that binds VPS26 at the N-terminus and VPS29 at the N-terminus. VPS35 also recruits the WASH complex by binding directly to the WASH complex subunit FAM21 (Harbour et al., 2012). WASH is a pentameric complex composed of WASH1, FAM21, CCDC53, strumpellin, and SWIP, and acts by promoting Arp2/3-dependent actin polymerization on endosomes. The coordinated action of WASH and Retromer link branched-actin formation with membrane tubulation, and promotes the scission of tubules containing the cargoes (Chen et al., 2019).

In non-neuronal cells, another major endosomal recycling pathway was recently discovered, which mediates the recycling from endosomes to the plasma membrane of over 120 cell surface proteins, including numerous cell adhesion proteins, signaling receptors and solute transporters (McNally et al., 2017). This pathway is mediated by a different member of the sorting nexin family of proteins, SNX17, and utilizes a multiprotein complex known as Retriever, which shares structural homology to the Retromer complex. The Retriever complex consists of three subunits: VPS35L, VPS26C and VPS29, the latter is shared with the Retromer complex. In addition, SNX17-dependent recycling requires the WASH complex and the CCC complex, which includes CCDC22 (coiled-coil domain containing 22)–CCDC93 (coiled-coil domain containing 93), and ten COMMD (copper metabolism MURR1 domain)-containing proteins (Chen et al., 2019; McNally and Cullen, 2018; Simonetti and Cullen, 2019; Wang et al., 2018a).

Prior to the discovery of the Retriever complex, SNX17 was shown to be present in neurons and regulate the trafficking of proteins including the amyloid precursor protein (APP) (Lee et al., 2008), LRP1 (Donoso et al., 2008) and ApoER2 (Sotelo et al., 2014; Feng et al., 2020). However, the roles and regulation of SNX17 and the Retriever at synaptic sites have not been directly tested.

Importantly, a growing body of evidence indicates that mutations in subunits of the SNX17-Retriever-CCC-WASH recycling pathway are involved in human pathologies primarily affecting the nervous system. For example, *VPS26C* is one of the genes that is overexpressed in Down syndrome (Lockstone et al., 2007). In addition, genetic defects in *VPS35L* and *CCDC22* have been linked to Ritscher-Schinzel syndrome, a disorder associated with developmental delay and intellectual disability (Kato et al., 2020; Kolanczyk et al., 2015; Voineagu et al., 2012; Otsuji et al., 2022). Moreover, mutations in *WASHC5*, which encodes the WASH complex subunit strumpellin, also underlie Ritscher-Schinzel syndrome (Elliott et al., 2013). In this case, mutations in WASH complex components would potentially impact both Retriever and Retromer recycling. However, despite these clear links with neurological disease, potential roles of SNX17 in synaptic function have not been characterized.

Among the best characterized cargoes of SNX17 is the adhesion molecule β1-integrin (Steinberg et al., 2012; Böttcher et al., 2012; McNally et al., 2017). Previous studies show that β1-integrin plays key roles in the regulation of synaptic function and plasticity. For example, immunogold labeling showed that β1-integrin localizes to synapses in CA1 hippocampal neurons, and is concentrated at the postsynaptic membrane (Mortillo et al., 2012). Pharmacological and genetic approaches altering the levels or function of β1-integrin or of integrin-associated kinases such as FAK and Src impair cytoskeletal organization at synapses and synaptic activity (Warren et al., 2012; Babayan et al., 2012; Orr et al., 2022). A postnatal forebrain and excitatory neuron-specific knockout of β1-integrin in the mouse showed impaired synaptic transmission through AMPA receptors as well as decreased NMDAR-dependent LTP, even though the steady-state expression of AMPAR subunits are not regulated by β1-integrin (Chan et al., 2006). Deletion of β1-integrin at postnatal stages impairs LTP (Huang et al., 2006), and interfering with β1-integrin function prevents the stabilization of LTP (Chun et al., 2001; Kramár et al., 2006). In addition, β1-integrin regulates the maturation of dendritic spines (Ning et al., 2013; Bourgin et al., 2007), and interfering with *α*5-integrin function, which forms heterodimers with β1-integrin, causes a reduction in the number of dendritic spines (Webb et al., 2007). Moreover, pharmacological activation of β1-integrin restored spine density and synaptic plasticity defects caused by loss of postsynaptic plasticity-related gene 1 (PRG-1) (Liu et al., 2016). In non-neuronal cells, β1-integrin trafficking through the endosomal pathway is a key regulatory step in the control of integrins at the cell surface. However, how β1-integrin recycling is regulated in neurons and importantly whether this regulation is coupled to synaptic activity has not been directly tested.

Here, we discover that the SNX17-Retriever pathway is critical for synapse function and synaptic plasticity. Specifically, we find that subunits of the SNX17 pathway partially co-localize with excitatory synapses. Moreover, we show that during long-term potentiation (LTP), the SNX17 pathway undergoes further recruitment to synapses. We discover that this recruitment is mediated by NMDAR-dependent Ca^2+^ influx, downstream CamKII signaling, and requires binding to the phosphoinositide, phosphatidylinositol 3-phosphate [PI(3)P] as well as the Retriever complex. Disrupting the SNX17 pathway induces loss of synapses and impairs structural changes in spines necessary for enduring changes in synapse function. Moreover, we show that these roles for the SNX17 pathway are due in part to SNX17-dependent control of surface levels of β1-integrin. Together, these findings reveal that SNX17 recycling is critical for dynamic changes at post-synaptic sites and plays a key role in regulating enduring forms of synaptic plasticity.

## Results

### The SNX17 pathway is present at synapses

SNX17-Retriever underlies a major recycling pathway in non-neuronal cells. Moreover, genetic alterations in proteins of the SNX17 recycling pathway are associated with several neurological disorders, which raises the possibility that SNX17 recycling may play a critical role in neuronal function. To better understand potential roles of SNX17 and Retriever in neurons, we used an antibody to detect endogenous SNX17 and analyzed the pattern of its expression in hippocampal neurons. DIV19 primary rat hippocampal neurons were fixed and stained for SNX17 and the dendritic marker MAP2. We found that SNX17 exhibited a punctate distribution pattern, with SNX17 puncta evident throughout the cell body and dendrites (Fig. 1A). We validated the specificity of the antibody by a knockdown approach, which resulted in reduced detection of SNX17-positive puncta (Fig. S1A). To examine SNX17 localization further, we tested if SNX17 is specifically present at excitatory synapses, which were detected by the overlap between the presynaptic marker vGLUT1 and the postsynaptic marker PSD95. We found that SNX17-positive puncta are present both at synaptic and extrasynaptic sites, with 44.6 ± 6.0% of structurally-defined synapses containing SNX17 (Fig. 1B and C). SNX17 showed similar colocalization with both PSD95 and vGLUT1 (Fig. 1B, D and E).

**Figure 1.**
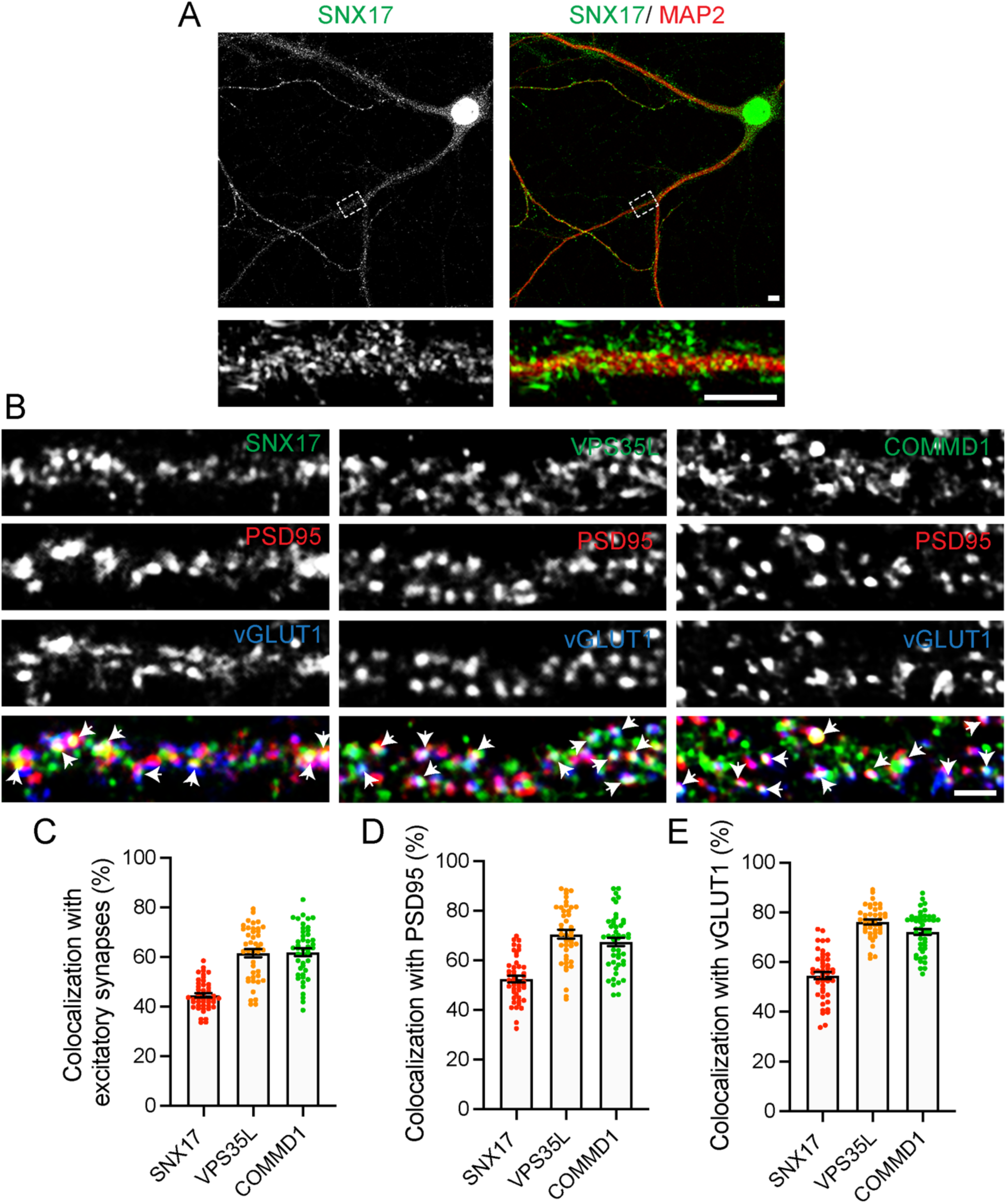
SNX17 is present in neurons and colocalizes with synaptic markers. (A) DIV19 rat hippocampal neurons were fixed and stained for SNX17 and the neuronal dendrite marker MAP2. Lower panels show straightened dendrites from the corresponding top panels. Scale bars, 2μm. (B) DIV19 rat hippocampal neurons were triple labelled with antibodies for the postsynaptic marker PSD95, the synaptic vesicle glutamate transporter vGLUT1 and either SNX17, VPS35L or COMMD1. Arrows indicate examples of colocalization of active excitatory synapses (labelled with both PSD95 and vGLUT1) with SNX17, VPS35L or COMMD1. Scale bar, 2μm. (C) The percentage of active excitatory synapses, labeled simultaneously with PSD95 and vGLUT1, that colocalize with the indicated proteins was determined using Mander’s colocalization coefficient (x100). SNX17: 44.600 ± 0.892, N=45 neurons; VPS35L: 61.610 ± 1.674, N=42 neurons; COMMD1: 61.980 ± 1.540, N=46 neurons. 3 independent experiments. Error bars are SEM. (D) The percentage of PSD95-labeled postsynaptic sites that colocalize with SNX17 was determined using Mander’s colocalization coefficient (x100). SNX17: 52.420 ± 1.336, N=45 neurons; VPS35L: 70.590 ± 1.816, N=42 neurons; COMMD1: 67.460 ± 1.690, N=46 neurons. 3 independent experiments. Error bars are SEM. (E) The percentage of vGLUT1-labeled presynaptic sites that colocalizes with SNX17 was determined using Mander’s colocalization coefficient (x100). SNX17: 54.600 ± 1.441, N=45 neurons; VPS35L: 76.150 ± 0.989, N=42 neurons; COMMD1: 72.150 ± 1.189, N=46 neurons. 3 independent experiments. Error bars are SEM.

The SNX17-dependent recycling pathway requires the participation of the Retriever, WASH and CCC complexes. Therefore, we tested if VPS35L, a core subunit of the Retriever, and COMMD1, a subunit of the CCC complex, are also localized to excitatory synapses. VPS35L and COMMD1 are, respectively, present in 61.6 ± 10.9% and 62.0 ± 10.3% of excitatory synapses defined by the co-localization of PSD95/vGLUT1 (Fig. 1B and C), and exhibit slightly higher colocalization with each synaptic marker individually (Fig. 1B, D and E). The abundance of SNX17 and other proteins of the recycling pathway at synapses suggests that SNX17-dependent recycling may play a role in regulating synaptic function.

### The SNX17 Retriever pathway is required to maintain excitatory synapses

To determine whether SNX17 is a regulator of synaptic strength, we used two different shRNA constructs to knockdown SNX17 in primary rat hippocampal neurons. Knockdown efficiency of the two shRNA clones was validated by lentiviral infection in the rat cell line Rat2 (Fig. S1B). We then transfected the shRNA constructs in rat hippocampal neurons and compared miniature excitatory postsynaptic currents (mEPSCs) of SNX17 shRNA-expressing neurons and scrambled shRNA control neurons. While mEPSC amplitude was not significantly altered by either SNX17 shRNA, the frequency of mEPSCs was significantly decreased (approximately 51.2% for clone 1 and 50.6% for clone 2) in SNX17 shRNA-expressing cells as compared with scrambled control shRNA-transfected neurons (Fig. 2A-C). We observed similar effects with both clones, and chose SNX17-shRNA clone 1 for all the following experiments. We further validated the knockdown efficiency of this clone by transduction in cortical neurons, which resulted in an 88.6 ± 0.9% decrease of SNX17 levels (Fig. S1C and D).

**Figure 2.**
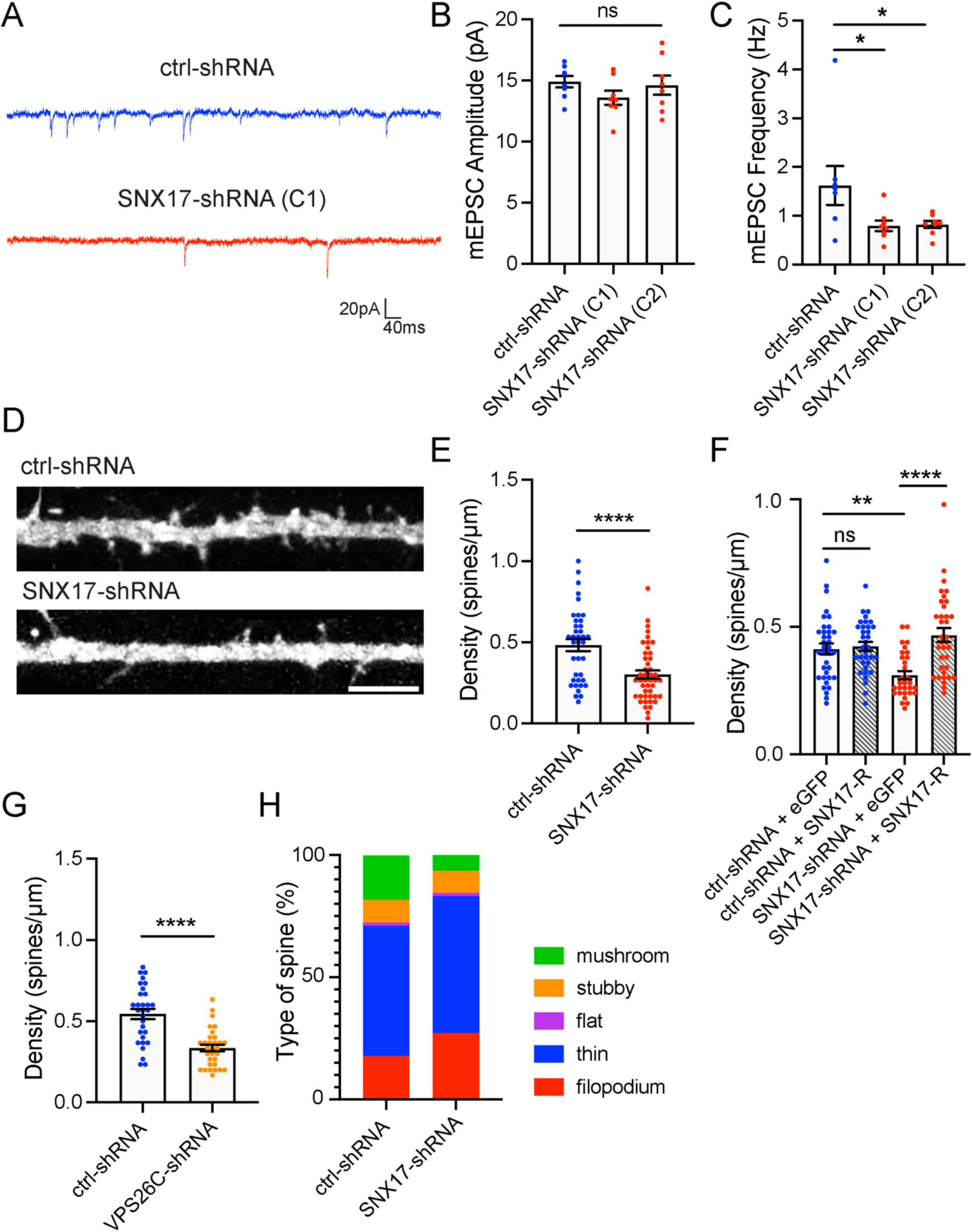
The SNX17 pathway regulates synaptic function by promoting dendritic spine maintenance. (A) Representative mEPSC recordings from DIV16-18 rat hippocampal neuron cultures transfected at DIV12 with a scramble control shRNA (ctrl-shRNA) or SNX17-shRNA clones, C1 or C2. (B) SNX17 knockdown with either shRNA clone did not significantly alter basal mEPSC amplitudes (one-way ANOVA). ctrl-shRNA: 14.910 ± 0.468, N=8 neurons; SNX17-shRNA C1: 13.600 ± 0.570, N=8 neurons; SNX17-shRNA C2: 14.630 ± 0.788, N=8 neurons. 3 independent experiments. Error bars are SEM. (C) SNX17 knockdown significantly decreased mEPSC frequency. ctrl-shRNA: 1.621 ± 0.398, N=8 neurons; SNX17-shRNA C1: 0.794 ± 0.109, N=8 neurons; SNX17-shRNA C2: 0.820 ± 0.073, N=8 neurons. Data were analyzed by one-way ANOVA with uncorrected Fisher’s LSD post hoc test, *p<0.05. Error bars are SEM. (D) Representative confocal images of dendritic spines in DIV16 hippocampal neurons co-transfected at DIV12 with eGFP (filler) and either ctrl-shRNA or SNX17-shRNA (C1). Scale bar, 5 µm. (E) Bar chart showing spine density (spines/μm). The number of dendritic spines in the first 30 μm of secondary dendrites was quantified. ctrl-shRNA: 0.483 ± 0.037, N=38 neurons; SNX17-shRNA: 0.302 ± 0.025, N=45 neurons. 3 independent experiments. Statistical significance was determined using unpaired two-tailed Student’s t-test, ****p<0.001. Error bars are SEM. (F) Bar chart showing spine density (spines/μm). DIV16 hippocampal neurons were co-transfected at DIV12 with mCherry (filler) and the indicated constructs. The numbers of dendritic spines in the first 50 µm of secondary dendrites were quantified. ctrl-shRNA + eGFP: 0.413 ± 0.021, N= 36 neurons; ctrl-shRNA + SNX17-R: 0.423 ± 0.018, N=34 neurons; SNX17-shRNA + eGFP: 0.310 ± 0.016, N=30 neurons; SNX17-shRNA + SNX17-R: 0.467 ± 0.027, N=35 neurons. Statistical significance was determined using one-way ANOVA with Tukey’s post hoc test, **p<0.01,****p<0.001. Error bars are SEM. (G) Hippocampal neurons were co-transfected at DIV12 with eGFP and either ctrl-shRNA or VPS26C-shRNA, fixed 4 days post-transfection, and the numbers of dendritic spines were quantified. ctrl-shRNA: 0.544 ± 0.032, N=30 neurons; VPS26C-shRNA: 0.334 ± 0.021, N=31 neurons. 3 independent experiments. Statistical significance was determined using unpaired two-tailed Student’s t-test, ****p<0.001. Error bars are SEM. (H) Descriptive graphic with the percentage of each spine type in ctrl-shRNA or SNX17-shRNA-transfected neurons. The dendritic spines in the first 30 μm of secondary dendrites were classified. ctrl-shRNA: N=30 neurons, SNX17-shRNA: N=30 neurons. 3 independent experiments.

A decrease in mEPSC frequency can be due to decreased release probability of presynaptic inputs or to a reduced number of synapses. To distinguish between these possibilities, we examined the effect of SNX17 knockdown on the density of dendritic spines, a structural hallmark of excitatory synaptic contacts. We transfected SNX17- and scramble control-shRNAs and, to ensure that we consistently sample spines from the same area for different neurons (McCartney et al., 2014), we quantified the number of spines in the first 50 μm of secondary dendrites. We found that SNX17 knockdown decreases the total number of dendritic spines by 37.5% (ctrl-shRNA: 0.483 ± 0.037 spines/μm, SNX17-shRNA: 0.302 ± 0.025 spines/μm) (Fig. 2D and E), similar to the reduction in mEPSC frequency. To evaluate whether loss of SNX17 accounts for these defects, we generated an shRNA-resistant GFP-SNX17 construct, and validated its resistance to knock-down in HEK293 cells (Fig. S1E). Transient transfection of shRNA-resistant GFP-SNX17 in cultured neurons rescued changes in spine density caused by SNX17 knockdown (Fig. 2F), which indicates that the knock-down-related defects are due to loss of SNX17. Moreover, knockdown of the core Retriever subunit VPS26C/DSCR3 caused a similar decrease in dendritic spine density (Fig. 2G).

We also evaluated whether SNX17 regulates dendritic spine morphology. We classified dendritic spines into the following classes: filopodial, thin, flat, stubby, and mushroom (Henry et al., 2017). Our results show that SNX17 knockdown causes a reduction in the number of filopodia and a decrease in the number of mushroom spines. In control shRNA-transfected neurons, the major spine type was thin (53.3 ± 2.5%), followed by mushroom (18.2 ± 2.2%), filopodial (17.8 ± 2.4%), stubby (9.5 ± 1.5%) and flat (1.1 ± 0.5%). In contrast, upon SNX17 knockdown, there was a striking 65% decrease in the number of mushroom-shaped spines, while filopodia became around 35% more abundant (thin: 56.0 ± 3.8%, mushroom: 6.4 ± 1.1%, filopodium: 27.2 ± 3.5%, stubby: 9.1 ± 2.1%, flat: 1.3 ± 0.4%) (Fig. 2H). Given that mushroom spines represent the most mature spine category (Hlushchenko et al., 2016), this data suggests that SNX17 plays a role in dendritic spine maturation. Together, these data suggest that the SNX17-Retriever recycling pathway regulates the density and maturation of excitatory synapses in hippocampal neurons.

### SNX17 is required for functional and structural plasticity upon cLTP

Given the changes in spine morphology, we tested whether the SNX17 pathway plays a role in enduring synaptic plasticity at excitatory synapses. We used a well-established chemical LTP (cLTP) induction protocol (400 µM glycine, 0 Mg^2+^; 5 min), which generates a long-lasting increase in postsynaptic strength (Lu et al., 2001; Park et al., 2004). As predicted, in neurons expressing the scrambled control shRNA, cLTP resulted in an increase in postsynaptic strength as assessed by an increase in mEPSC amplitude and frequency in whole-cell patch-clamp recordings. Notably, SNX17 knockdown blocked the cLTP-induced increase in mEPSC amplitude and frequency (Fig. 3A-C). This effect was observed in mEPSCs recorded in the first 30 min after cLTP, as well as at later time points, which indicates that SNX17 knockdown blocks the initiation of cLTP (Fig. S1F and G).

**Figure 3.**
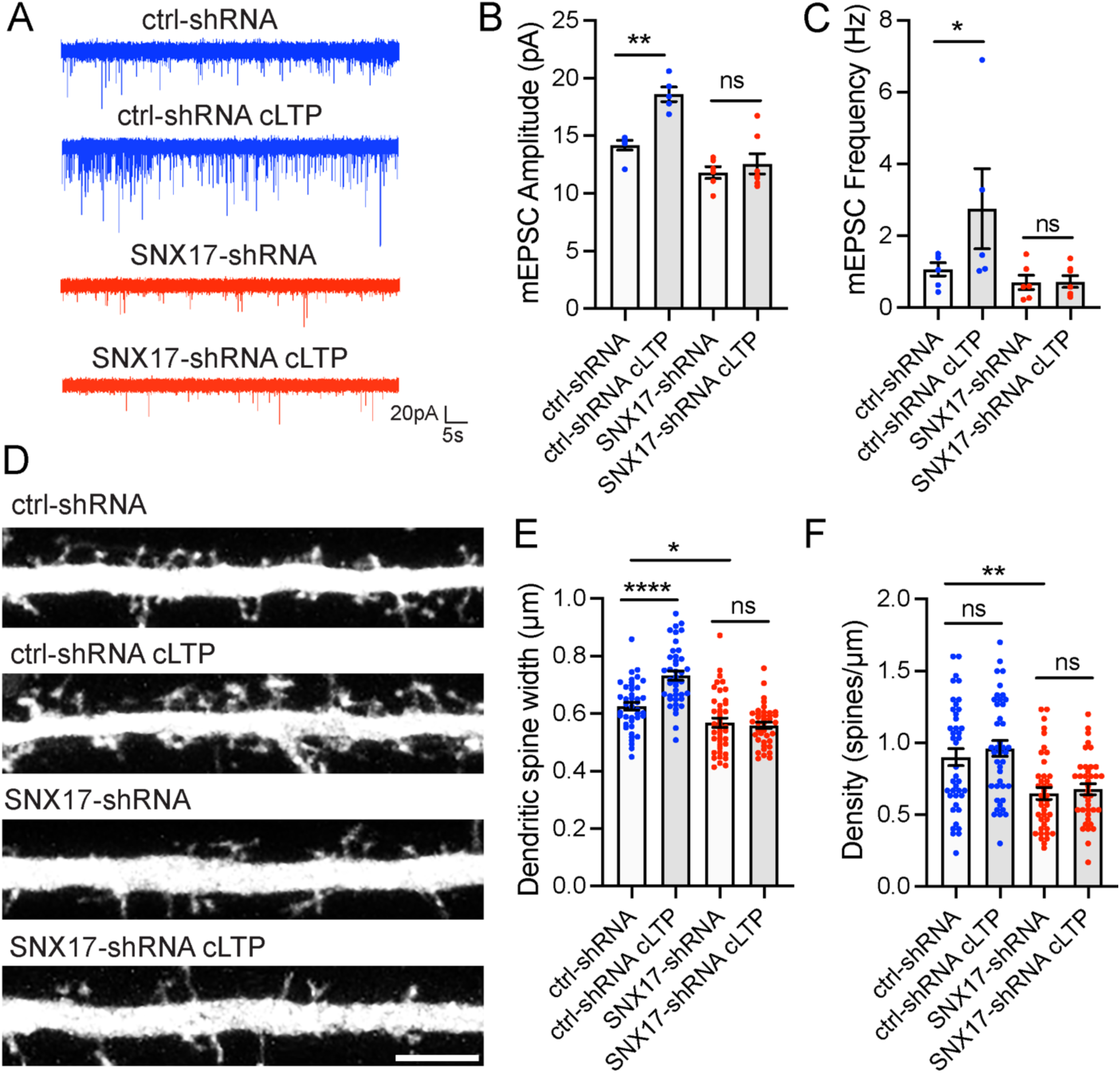
SNX17 is required for functional and structural plasticity during cLTP. (A) Example traces for DIV16-18 rat hippocampal neuron cultures transfected at DIV12 with a scramble control shRNA (ctrl-shRNA) or SNX17-shRNA, and either treated with cLTP or left untreated. (B) Induction of cLTP increased the amplitude of mEPSCs in neurons transfected with ctrl-shRNA but not SNX17-shRNA. ctrl-shRNA: 14.190 ± 0.426, N=6 neurons; ctrl-shRNA with cLTP: 18.610 ± 0.638, N=5 neurons; SNX17-shRNA: 11.820 ± 0.505, N=6 neurons; SNX17-shRNA with cLTP: 12.580 ± 0.881, N=7 neurons. 3 independent experiments. Data were analyzed by one-way ANOVA with uncorrected Fisher’s LSD post hoc test, **p<0.01. Error bars are SEM. (C) Induction of cLTP increased the frequency of mEPSCs in neurons transfected with ctrl-shRNA but not SNX17-shRNA. ctrl-shRNA: 1.069 ± 0.182, N=6 neurons; ctrl-shRNA with cLTP: 2.751 ± 1.116, N=5 neurons; SNX17-shRNA: 0.705 ± 0.200, N=6 neurons; SNX17-shRNA with cLTP: 0.726 ± 0.162, N=7 neurons. 3 independent experiments. Data were analyzed by one-way ANOVA with uncorrected Fisher’s LSD post hoc test, *p<0.05. Error bars are SEM. (D) Representative confocal images of dendritic spines in DIV16 hippocampal neurons co-transfected at DIV12 with eGFP (filler) and either ctrl-shRNA or SNX17-shRNA. Neurons were either treated with cLTP or left untreated, and fixed 50 min after cLTP. Scale bar, 5 µm. (E) The maximum width for each spine was quantified, and the average size of the dendritic spines in the first 30 μm of secondary dendrites was calculated. ctrl-shRNA: 0.625 ± 0.013, N=41 neurons; ctrl-shRNA with cLTP: 0.732 ± 0.016, N=41 neurons; SNX17-shRNA: 0.568 ± 0.016, N=41 neurons; SNX17-shRNA with cLTP: 0.559 ± 0.011, N=38 neurons. 3 independent experiments. Data were analyzed by one-way ANOVA with Tukey’s post hoc test, *p<0.05, ****p<0.001. Error bars are SEM. (F) Quantification of spine density (spines/μm). ctrl-shRNA: 0.900 ± 0.058, N=41 neurons; ctrl-shRNA with cLTP: 0.961 ± 0.054, N=41 neurons; SNX17-shRNA: 0.647 ± 0.042, N=41 neurons; SNX17-shRNA with cLTP: 0.677 ± 0.038, N=38 neurons. 3 independent experiments. Data were analyzed by one-way ANOVA with Tukey’s post hoc test, **p<0.01. Error bars are SEM.

LTP is associated with increased expression of AMPA-type glutamate receptors at synapses, as well as structural increases in the spine head area. These changes often appear together, although some studies demonstrate that they are regulated by distinct molecular pathways (Nakahata and Yasuda, 2018; Citri and Malenka, 2008). To determine if the SNX17-Retriever pathway regulates structural plasticity of dendritic spines, we quantified spine head width in the first 30 μm of secondary dendrites in SNX17 knockdown neurons treated with or without cLTP stimulation. Spine head enlargement can be observed as early as 15 minutes post-glycine stimulation, but it continues to increase over time and is more evident 45 minutes after glycine stimulation (Henry et al., 2017). As expected, cells expressing a scrambled (control) shRNA demonstrated a significant increase in spine head area 50 minutes after cLTP induction, but this structural plasticity was lost in SNX17 knockdown neurons (Fig. 3D and E). We did not observe overall changes in dendritic spine density in control neurons following cLTP (Fig. 3F). Similarly, while spine density was lower in SNX17 knockdown neurons, this lower spine density remained constant following cLTP (Fig. 3F). In addition, shRNA-mediated knockdown of the Retriever subunit VPS26C (Fig. S2A), similarly blocked the increase in dendritic spine size following cLTP (Fig. S2 B-D). Together, these data indicate that the SNX17-Retriever recycling pathway is critical for the structural and functional synaptic changes that underlie enduring enhancement of synapse function following cLTP.

### SNX17 is recruited to excitatory synapses during cLTP, and this recruitment is dependent on neuron-specific CamKII signaling

cLTP is characterized by extensive synapse remodeling and changes in surface-exposed channels and receptors (van Oostrum et al., 2020). These regulatory changes occur via the controlled delivery of membrane proteins from intracellular compartments, although the contribution of the SNX17-Retriever recycling pathway to this process is unknown. We therefore investigated whether SNX17-dependent recycling is engaged during cLTP.

Under basal conditions, there is a prominent synaptic localization of SNX17 and other SNX17-Retriever pathway subunits (Fig. 1). In addition, SNX17 is also found at extrasynaptic sites. We therefore asked whether the patterned synaptic activity that drives LTP impacts the localization of SNX17 at synaptic sites. We induced cLTP in DIV19 cultured hippocampal neurons and using immunocytochemistry, analyzed the relative synaptic localization of SNX17 at different time points after cLTP. Importantly, there was around 19.4% and 15.2% increase, respectively, in the co-localization of SNX17 with the excitatory synaptic markers vGLUT1 and PSD95 at 10 and 30 min post-cLTP. At 60 min post-cLTP, the synaptic localization of SNX17 returned to basal levels (baseline: 43.310 ± 1.881%, 10 min after cLTP: 51.750 ± 1.520%, 30 min after cLTP: 49.930 ± 1.533%, 60 min after cLTP: 39.680 ± 1.625%) (Fig. 4A and B). We also transfected rat hippocampal neurons with eGFP as a filler and quantified the intensity of SNX17 at dendritic spines in cells treated with or without cLTP. We found a similar 20% increase in the intensity of SNX17 at dendritic spines 10 minutes after LTP (mean intensity for baseline: 0.505 ± 0.022, mean intensity for cLTP: 0.602 ± 0.024) (Fig. 4C). In addition, the synaptic localization of the core Retriever subunit VPS35L followed a similar trend, with an approximate 19% increase in its localization to excitatory synapses 10 min post-LTP and a return to baseline levels after 60 min (baseline: 58.330 ± 1.725%, 10 min after cLTP: 69.380 ± 1.917%, 30 min after cLTP: 65.730 ± 1.898%, 60 min after cLTP: 56.380 ± 2.228% (Fig. S2E and F).

**Figure 4.**
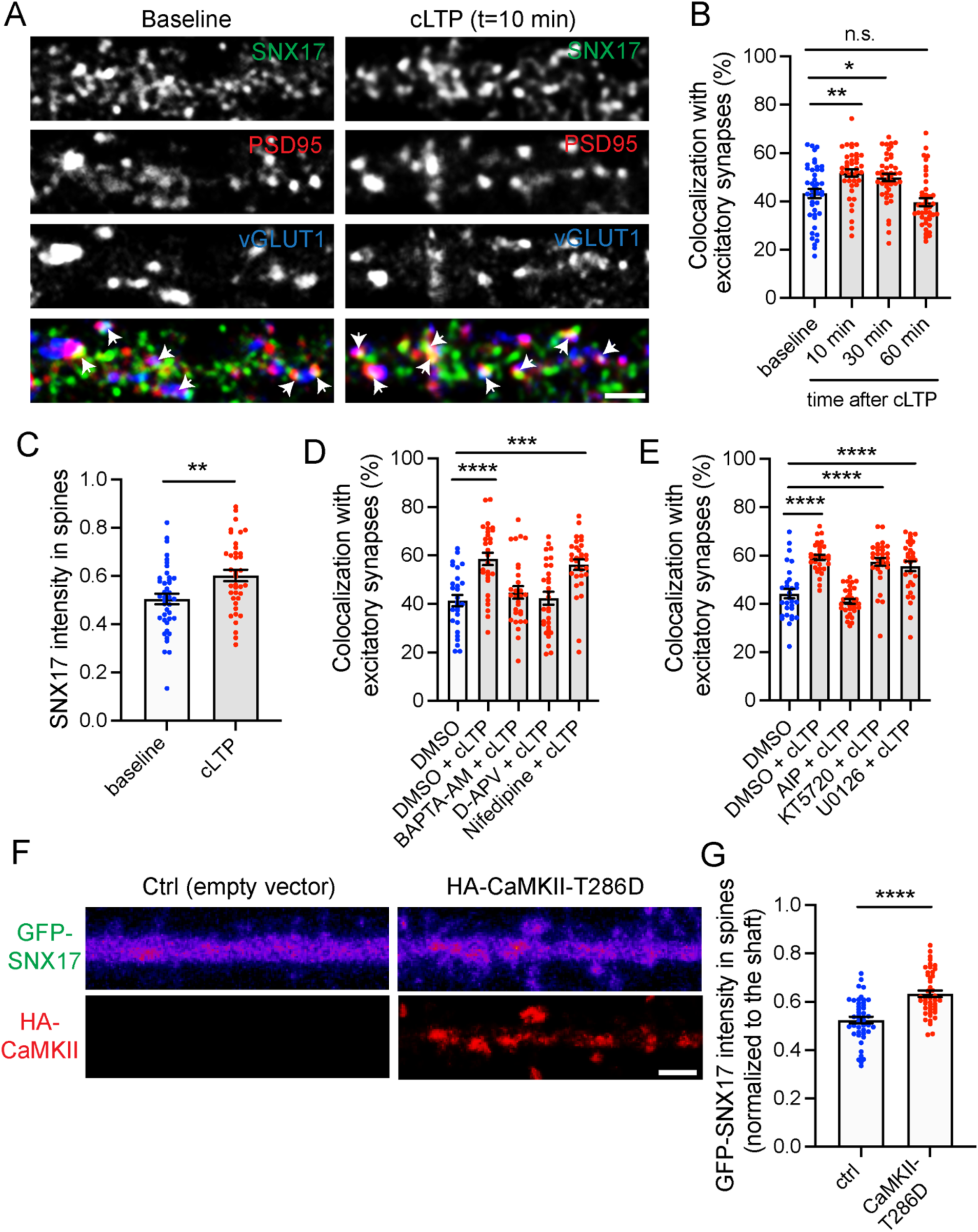
SNX17 is dynamically recruited to synapses during cLTP, which is dependent on the NMDAR-Calcium-CaMKII pathway. (A) Representative confocal images showing the colocalization of SNX17 with excitatory synapses before and 10 min after cLTP treatment. DIV19 rat hippocampal neuron cultures were untreated or treated with cLTP for 5 min (Mg^2+^−free HBS with: 400 μM glycine, 20 μM bicuculline, and 3 μM strychnine). Cells were fixed after cLTP at 10, 30, or 60 min, permeabilized and incubated with antibodies against SNX17, PSD95 and vGLUT1. Arrows indicate example of colocalization. Scale bar, 2 µm. (B) The percentage of excitatory synapses that colocalize with SNX17 at the different time points was determined using Mander’s colocalization coefficient (x100). Baseline: 43.310 ± 1.881, N=43; 10 min after cLTP: 51.750 ± 1.520, N=42; 30 min after cLTP: 49.930 ± 1.533, N=43; 60 min after cLTP: 39.680 ± 1.625, N=43. 3 independent experiments. Data were analyzed by one-way ANOVA with Tukey’s post hoc test, *p<0.05, **p<0.01. Error bars are SEM. (C) Hippocampal neurons were transfected at DIV12 with eGFP, and 4 days post-transfection either treated with cLTP for 5 min or left untreated. Cells were washed and further incubated with HBS for 10 min, followed by fixation and immunocytochemistry with an anti-SNX17 antibody. The mean intensity of SNX17 in the spines present in the first 30 µm of secondary dendrites was quantified and normalized to SNX17 mean intensity in the dendritic shaft. Baseline: 0.505 ± 0.022, N=41 neurons; cLTP: 0.602 ± 0.024, N=39 neurons. Statistical significance was determined using unpaired two-tailed Student’s t-test, **p<0.01. Error bars are SEM. (D) DIV19 rat hippocampal neuron cultures were treated with DMSO, 10 μM BAPTA-AM, 100 μM D-APV or 10 μM nifedipine for 30 min, followed by a 5-min cLTP stimulus in the presence of compounds where indicated. Neurons were further incubated in the presence of the indicated compounds for 10 min before fixation, followed by permeabilization and incubation with antibodies against SNX17, PSD95 and vGLUT1. The percentage of excitatory synapses that colocalize with SNX17 at the different time points was determined using Mander’s colocalization coefficient (x100). DMSO: 41.380 ± 2.328, N=28 neurons; DMSO + cLTP: 58.550 ± 2.538, N=30 neurons; BAPTA-AM + cLTP: 44.710 ± 2.597, N=29 neurons; D-APV + cLTP: 42.370 ± 2.631, N=30 neurons; nifedipine + cLTP: 56.190 ± 2.150, N=31 neurons. 3 independent experiments. Data were analyzed by one-way ANOVA with Tukey’s post hoc test, ***p<0.005, **** p<0.001. Error bars are SEM. (E) DIV19 rat hippocampal neurons were treated with DMSO, 10 μM AIP, 2 μM KT5720, or 10 μM U0126 for 30 min, followed by a 5-min cLTP stimulus in the presence of compounds where indicated. Neurons were further incubated in the presence of the indicated compounds for 10 min before fixation, followed by permeabilization and incubation with antibodies against SNX17, PSD95 and vGLUT1. The percentage of excitatory synapses that colocalize with SNX17 at the different time points was determined using Mander’s colocalization coefficient (x100). DMSO: 44.250 ± 1.951, N=30 neurons; DMSO + cLTP: 59.220 ± 1.124, N=30 neurons; AIP + cLTP: 41.090 ± 1.002, N=31 neurons; KT5720 + cLTP: 57.400 ± 1.654, N=31 neurons; U0126 + cLTP: 55.590 ± 2.015, N=30 neurons. 3 independent experiments. Data were analyzed by one-way ANOVA with Tukey’s post hoc test, ***p<0.005, **** p<0.001. Error bars are SEM. (F) Representative confocal images of DIV17 hippocampal neurons co-transfected at DIV16 with GFP-SNX17 and either an empty pRK5 vector or pRK5-HA-CaMKII-T286D. Neurons were fixed, permeabilized and incubated with an anti-HA antibody. GFP-SNX17 intensity presented in the “fire” LUT color scheme. Scale bar, 2 µm. (G) The mean intensity of GFP-SNX17 was measured in the dendritic spines present in the first 30 μm of secondary dendrites, and normalized to GFP-SNX17 mean intensity in the dendritic shaft. Ctrl: 0.525 ± 0.014, N=45 neurons; CaMKII-T286D: 0.633 ± 0.014, N=45 neurons. Statistical significance was determined using unpaired two-tailed Student’s t-test, **** p<0.001. Error bars are SEM.

cLTP requires N-methyl-D-aspartate (NMDA) receptor activation, which results in an increase in postsynaptic calcium and the activation of several downstream signaling cascades, which in turn result in an increase in synaptic efficacy (Musleh et al., 1997). Therefore, we investigated whether the cLTP-dependent SNX17 recruitment to synapses was mediated by NMDA receptor activation and an increase in intracellular calcium. We chose three manipulations: chelating intracellular calcium with membrane-permeable BAPTA-AM, blocking NMDA receptors with D-APV treatment, or blocking L-type voltage-dependent calcium channels (VDCCs) with nifedipine. We analyzed SNX17 recruitment to synapses 10 min after cLTP stimulus, because this time point showed the maximum colocalization with excitatory synapses under basal conditions. BAPTA-AM and D-APV treatment completely blocked the cLTP-dependent increase in SNX17 localization to excitatory synapses. Surprisingly, despite the fact that VDCCs are responsible for around 80% of the total calcium entry in response to presynaptic glutamate release (Schiller et al., 1998), nifedipine treatment had no effect on SNX17 recruitment (Fig. 4D). This data suggests that initial calcium entry through glycine-stimulated NMDA receptors is primarily responsible for driving SNX17 recruitment to synapses.

During LTP, many signaling enzymes are activated including calcium/calmodulin-dependent kinase II (CamKII), protein kinase A (PKA) and Ras-extracellular signal-regulated kinase (ERK) (Lisman et al., 2012). Calcium entry activates these kinases, which in turn activate signaling pathways that result in increased synaptic transmission. We found that AIP, a selective and potent inhibitor of CamKII (Ishida et al., 1995), blocked SNX17 recruitment to synapses upon cLTP stimulation. However, both KT5720 and U0126, which inhibit PKA- and ERK-dependent signaling pathways, respectively, failed to block cLTP-dependent SNX17 recruitment (Fig. 4E).

Based on previous studies, addition of an N-terminal tag to SNX17 does not interfere with its binding to cargo or endosomes (McNally et al., 2017). Indeed, we found by western blot of transfected HEK293 cells that GFP-SNX17 is expressed (Fig. S3A), and in rat hippocampal neurons, GFP-SNX17 localizes to EEA1-positive endosomes and to PSD95-positive synapses (Fig. S3B and C).

To test whether CaMKII activation is sufficient to drive SNX17 recruitment to synapses, we utilized a CaMKII mutant, T286D, which increases CaMKII activity via mimicking phosphorylation at a site of autophosphorylation (Fong et al., 1989). This mutant induces LTP when introduced with a viral expression system (Pettit et al., 1994) or by direct injection into postsynaptic cells (Lledo et al., 1995). We co-transfected rat hippocampal neurons with CaMKII-T286D (or an empty vector) together with a GFP-SNX17 construct, and found that CaMKII-T286D expression for 24 h drove GFP-SNX17 recruitment to dendritic spines in the absence of an external cLTP stimulus (mean intensity for ctrl-empty vector: 0.53 ± 0.01, mean intensity for CaMKII-T286D: 0.63 ± 0.01) (Fig. 4F and G). These findings indicate that CaMKII activation is sufficient to promote SNX17 recruitment to synapses. Together, our results indicate that cLTP stimulation induces a transient recruitment of SNX17 and other SNX17-Retriever pathway subunits to excitatory synapses and that activity-dependent CaMKII-signaling is upstream of this SNX17-recruitment.

To monitor the dynamic recruitment of SNX17 and test whether there is a correlation with the cLTP-dependent structural changes at individual spines, we performed live-cell imaging of GFP-SNX17 in neurons co-expressing soluble mCherry (Fig. 5A). Importantly, cLTP resulted in an increase in spine head area 30 min after cLTP treatment (Fig. 5B). Similar to endogenous SNX17 (Fig. 4B), we observed maximum recruitment to synapses 10 min following cLTP induction, with a 31.6 ± 0.05 % increase in GFP-SNX17 intensity at dendritic spines as compared to baseline intensity. GFP-SNX17 intensity at dendritic spines remained significantly elevated during the remaining time points analyzed, up to 30 min after the cLTP stimulus (Fig. 5C). Analysis of these live cell imaging studies revealed a significant correlation between the increase in GFP-SNX17 intensity in spines at t=10 min and the growth of individual spines 30 min post-cLTP (Fig. 5D). These results indicate that SNX17 is actively recruited to synapses during cLTP and this recruitment is specifically related to structural enlargement of spines necessary for enduring increases in synaptic function.

**Figure 5.**
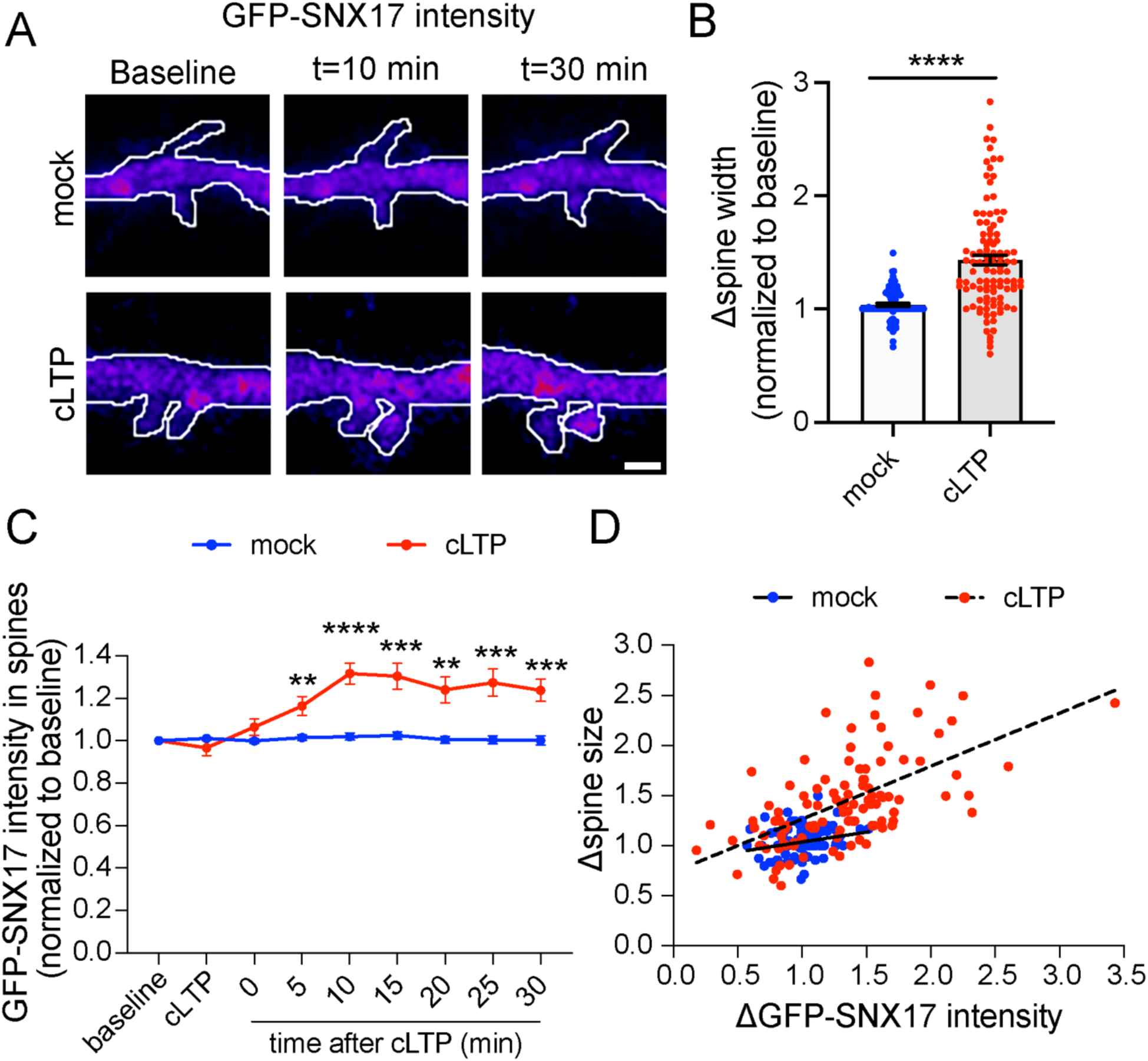
The cLTP-dependent recruitment of SNX17 to dendritic spines correlates with increased dendritic spine width. (A) Example confocal images taken from DIV17 neurons expressing GFP-SNX17 and mCherry (filler) under conditions of cLTP or HBS control (mock). Images of live neurons in an environmental chamber were captured before cLTP (baseline), during cLTP, and at 0, 5, 10, 15, 20, 25 and 30 min after the cLTP stimulus. Representative images of baseline, t=10 and t=30 are shown. Intensity presented in the “fire” LUT color scheme. Scale bar, 1 µm. (B) The endpoint of the experiment (30 min) was used to measure the cLTP-dependent increase in spine width. Spine width was normalized to the baseline size for each spine. Mock: 1.038 ± 0.014, N=95 spines across 12 cells; cLTP: 1.432 ± 0.044, N=106 spines across 12 cells. Statistical significance was determined using unpaired two-tailed Student’s t-test, ****p<0.001. Error bars are SEM. (C) The mean intensity of GFP-SNX17 was measured in the spines that remained in the same plane at each time point following cLTP (or mock) and normalized to the baseline for each individual spine. Mock: N=95 spines across 12 cells (baseline: 1.000, cLTP: 1.011 ± 0.008, 0: 0.999 ± 0.011, 5: 1.014 ± 0.014, 10: 1.020 ± 0.017, 15: 1.025 ± 0.017, 20: 1.005 ± 0.017, 25: 1.004 ± 0.019, 30: 1.001 ± 0.021); cLTP: N=106 spines across 12 cells (baseline: 1.000, cLTP: 0.966 ± 0.035, 0: 1.064 ± 0.038, 5: 1.163 ± 0.045, 10: 1.316 ± 0.049, 15: 1.304 ± 0.061, 20: 1.239 ± 0.061, 25: 1.274 ± 0.063, 30: 1.237 ± 0.052). Statistical significance was determined using 2-way ANOVA with Sidak’s multiple comparison test, **p<0.01, ***p<0.005, ****p<0.001. Error bars are SEM. (D) Scatter plot of individual changes in spine width at the endpoint of the experiment vs changes in GFP-SNX17 intensity at 10 min. Mock: N=95 spines across 12 cells, cLTP: N=106 spines across 12 cells. A linear regression model was fit to the cLTP (R squared=0.3483, p<0.0001) and mock (R squared=0.0573, p=0.0195) conditions.

To determine the relationship between synaptic recruitment of SNX17 and its association with the Retriever complex, we examined the dynamic localization of an L470G mutant of GFP-SNX17, which is defective in binding to Retriever (McNally et al., 2017). This mutant was generated on top of the shRNA-resistant version of GFP-SNX17. It had similar expression levels to WT and shRNA-resistant GFP-SNX17 (Fig. S3A), exhibited a punctate intracellular localization, and maintained colocalization with EEA1 (Fig. S3D). Notably the L470G mutant failed to colocalize with PSD95-positive synapses (Fig. S3E). However, neuronal expression of the L470G SNX17 mutant did not block the cLTP-stimulated increase in dendritic spine width, presumably due to the presence of endogenous SNX17 (Fig. 6A and B). Dynamic imaging studies revealed that unlike WT GFP-SNX17 (Fig. 5C), cLTP stimulation fails to drive synaptic recruitment of the L470G SNX17 mutant (Fig. 6C). These results demonstrate that in order for its recruitment to synapses, SNX17 must be bound to the Retriever complex.

**Figure 6.**
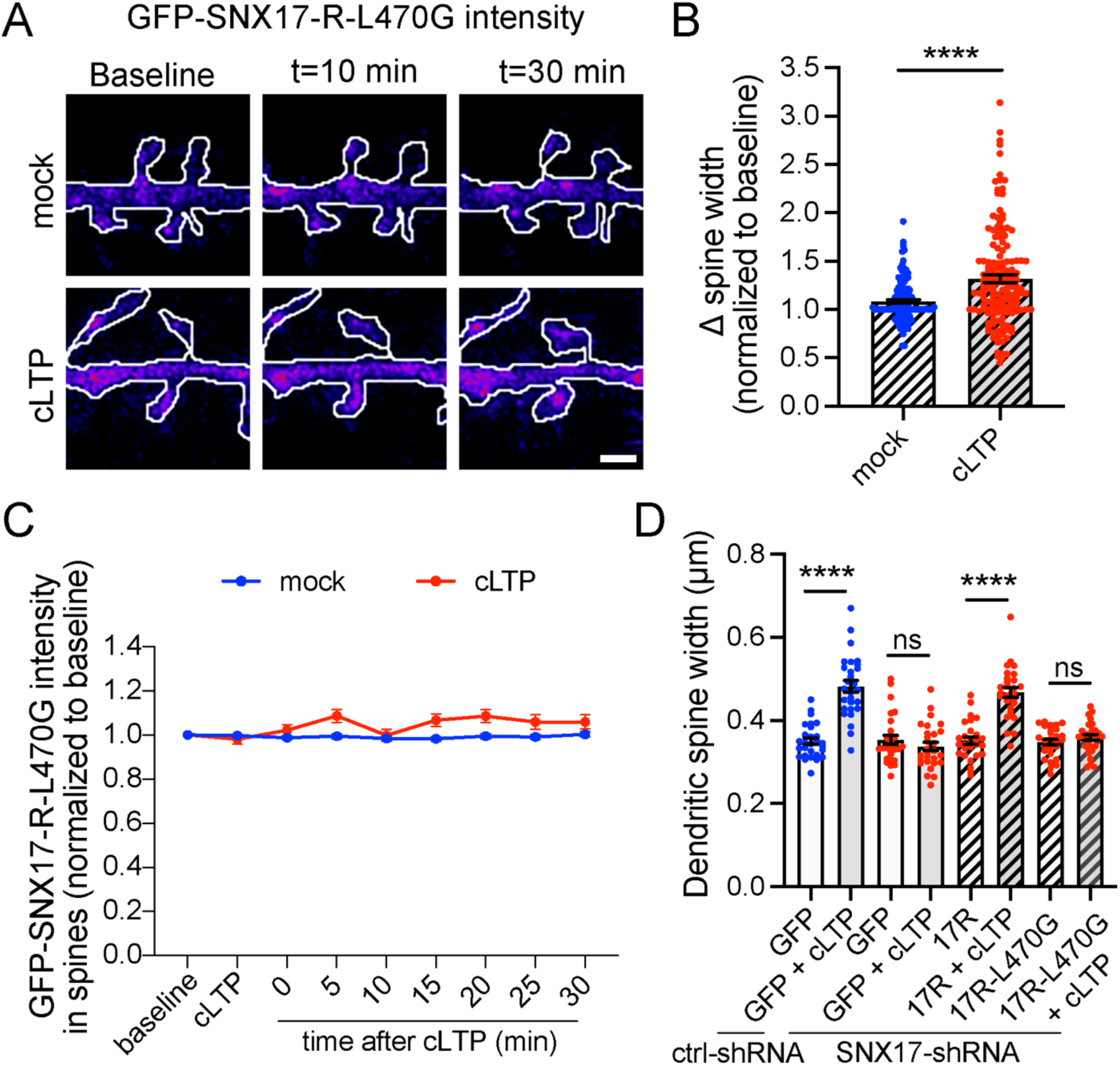
Retriever binding is necessary for the cLTP-dependent recruitment of SNX17 to dendritic spines as well as the for SNX17-mediated increase in dendritic spine width following cLTP. (A) Example confocal images taken from DIV17 neurons expressing GFP-SNX17-R-L470G and mCherry (filler) under conditions of cLTP or HBS control (mock). Images of live neurons in an environmental chamber were captured before cLTP (baseline), during cLTP, and at 0, 5, 10, 15, 20, 25 and 30 min after the cLTP stimulus. Representative images of baseline, t=10 and t=30 are shown. Intensity presented in the “fire” LUT color scheme. Scale bar, 1 µm. (B) The endpoint of the experiment (30 min) was used to measure the cLTP-dependent increase in spine width. Spine width was normalized to the baseline size for each spine. Mock: 1.081 ± 0.020, N=95 spines across 12 cells; cLTP: 1.318 ± 0.039, N=146 spines across 12 cells. Statistical significance was determined using unpaired two-tailed Student’s t-test, ****p<0.001. Error bars are SEM. (C) The mean intensity of GFP-SNX17-R-L470G was measured in the spines that remained in the same plane at each time point following cLTP (or mock) and normalized to the baseline for each individual spine. Mock: N=95 spines across 12 cells (baseline: 1.000, cLTP: 0.997 ± 0.006, 0: 0.988 ± 0.009, 5: 0.994 ± 0.011, 10: 0.984 ± 0.010, 15: 0.982 ± 0.010, 20: 0.995 ± 0.012, 25: 0.991 ± 0.012, 30: 1.003 ± 0.012); cLTP: N=146 spines across 12 cells (baseline: 1.000, cLTP: 0.980 ± 0.020, 0: 1.022 ± 0.025, 5: 1.086 ± 0.030, 10: 0.999 ± 0.027, 15: 1.067 ± 0.029, 20: 1.086 ± 0.030, 25: 1.059 ± 0.033, 30: 1.059 ± 0.032). Two-way ANOVA with Sidak’s multiple comparison test determined that there are no significant differences between the mock and cLTP conditions. Error bars are SEM. (D) DIV16 hippocampal neurons were co-transfected at DIV12 with mCherry (filler) and the indicated constructs. Neurons were either treated with cLTP or left untreated, and fixed 50 min after cLTP. The maximum width for each spine was quantified, and the average size of the dendritic spines in the first 30 μm of secondary dendrites was calculated. ctrl-shRNA + GFP: 0.350 ± 0.008, N=28 neurons; ctrl-shRNA + GFP + cLTP 0.482 ± 0.014, N=28 neurons; SNX17-shRNA + GFP: 0.353 ± 0.011, N=27 neurons; SNX17-shRNA + GFP + cLTP: 0.338 ± 0.010, N=28 neurons; SNX17-shRNA + 17R: 0.352 ± 0.009, N=28 neurons; SNX17-shRNA + 17R + cLTP: 0.468 ± 0.012, N=28 neurons; SNX17-shRNA + 17R-L470G: 0.348 ± 0.007, N=27 neurons; SNX17-shRNA + 17R-L470G + cLTP: 0.359 ± 0.007, N=28 neurons. 3 independent experiments. Data were analyzed by one-way ANOVA with Tukey’s post hoc test, ****p<0.001. Error bars are SEM.

Given that SNX17 recruitment to dendritic spines requires that it bind to Retriever, we investigated whether structural remodeling of dendritic spines during cLTP requires the association of SNX17 with Retriever. We depleted endogenous SNX17 with shRNA, then tested the ability of WT SNX17 or the L470G SNX17 mutant to rescue spine enlargement after cLTP induction. Indeed, shRNA-resistant GFP-SNX17 fully rescued cLTP-induced changes in spine head width to levels similar to controls, however there was no significant rescue by shRNA-resistant L470G GFP-SNX17 (Fig. 6D). These studies indicate that Retriever binding to SNX17 is necessary for SNX17 recruitment to synapses as well as for cLTP-dependent changes at synapses that occur downstream of SNX17 recruitment.

In addition to GFP-SNX17 recruitment to dendritic spines, we also observed extrasynaptic changes in the formation of SNX17-positive puncta in dendritic shafts following cLTP (Fig. 7A). The number of GFP-SNX17 puncta increased gradually in cLTP-treated neurons, but remained constant in mock-treated cells. Significant differences first appeared at t=10 min, with a 1.5 fold increase compared to baseline, and became 2-fold higher at t=30 min (Fig. 7B). However, there were no significant changes in puncta size at the time points analyzed (Fig. 7C). Retriever binding is necessary for the cLTP-dependent increase in the number of SNX17-positive puncta, as the L470G mutant failed to show an increase in puncta numbers following cLTP (Fig. 7D). One possibility is that the increased numbers of puncta reflect increased expression of SNX17-Retriever pathway subunits via upregulation of protein synthesis. However, we did not detect changes in the total protein levels of SNX17, VPS35L and COMMD1 following cLTP (Fig. S4 A-D). Alternatively, cLTP may promote the recruitment of cytosolic SNX17 to endosomal compartments. To test this possibility, we quantified the localization of SNX17 with the endosomal markers EEA1 and VPS35. EEA1 is a marker for early endosomes, and extensively colocalizes with SNX17 in an epithelial cell line (Steinberg et al., 2012). VPS35 is a marker of the Retromer complex, and both Retromer and Retriever pathways emerge from the same microdomains of VPS35-containing endosomes (McNally et al., 2017; Singla et al., 2019). Intriguingly, after 10 min of a cLTP stimulus, we found an increase in the co-localization of SNX17 with VPS35 (Fig. 7E and F) with no changes in the co-localization with EEA1 (Fig. S4E and F). These results suggest that during cLTP, SNX17 is recruited to endosomal compartments that are active in recycling. As an additional marker for endosomes that are active in recycling we used Syntaxin 13 (Prekeris et al., 1998), and found an increase in the colocalization of SNX17 with Syntaxin 13 following cLTP (Fig. 7G).

**Figure 7.**
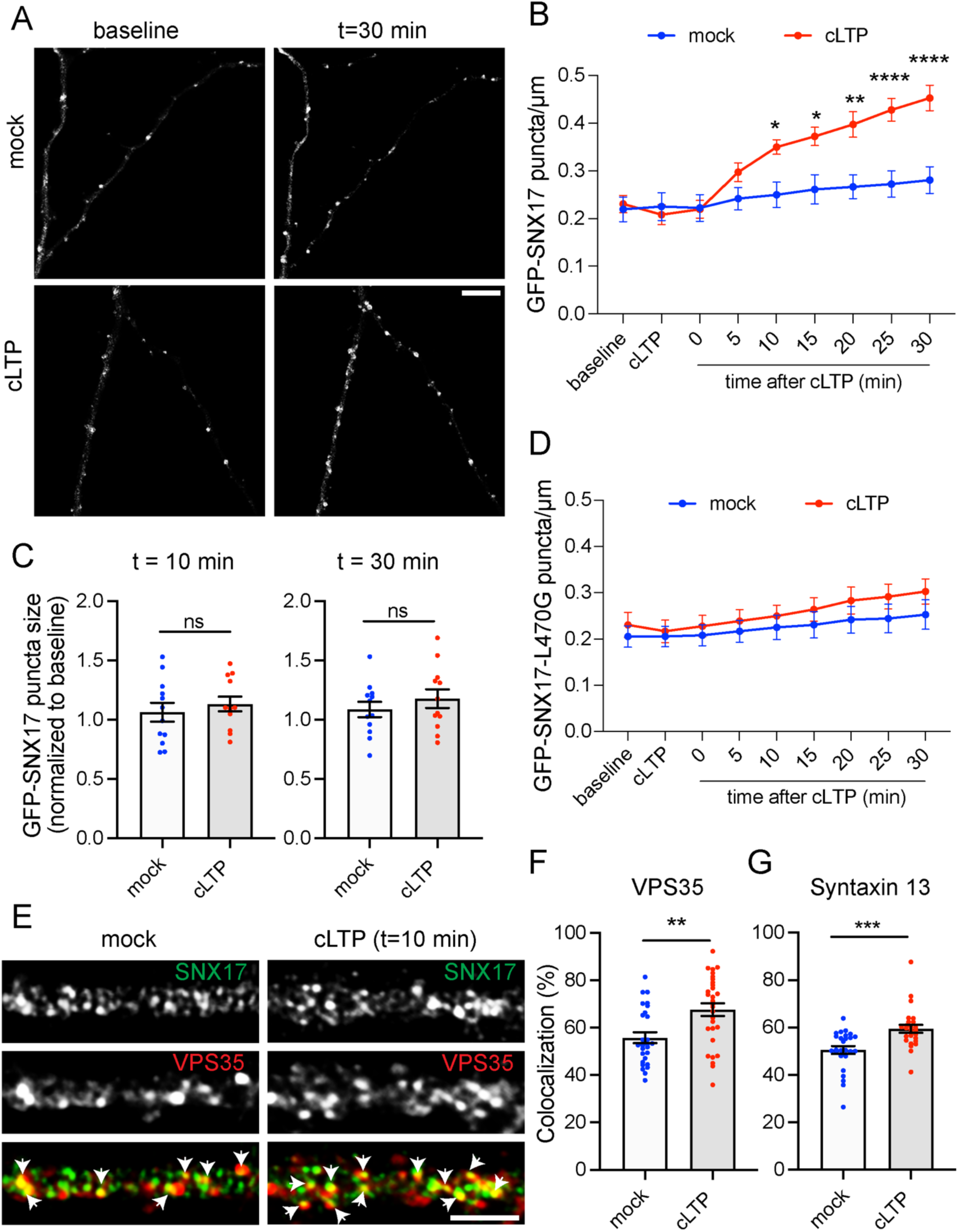
cLTP promotes extrasynaptic changes in the formation of SNX17-positive puncta, which may correspond to endosomal compartments that are active in recycling. (A) Example confocal images of secondary dendrites from DIV17 neurons expressing GFP-SNX17 and mCherry (filler) under conditions of cLTP or HBS control (mock). Images of live neurons in an environmental chamber were captured before cLTP (baseline), during cLTP, and at 0, 5, 10, 15, 20, 25 and 30 min after the cLTP stimulus. Representative images of baseline and t=30 are shown. Scale bar, 5 µm. (B) The number of GFP-SNX17-positive puncta in 30 µm of dendritic spines at the different time points following cLTP (or mock) was quantified and normalized to the baseline for each dendrite. Mock: N=12 dendrites (baseline: 0.219 ± 0.026, cLTP: 0.225 ± 0.029, 0: 0.222 ± 0.028, 5: 0.242 ± 0.023, 10: 0.250 ± 0.026, 15: 0.261 ± 0.031, 20: 0.267 ± 0.025, 25: 0.272 ± 0.027, 30: 0.281 ± 0.028); cLTP: N=12 dendrites (baseline: 0.231 ± 0.018, cLTP: 0.208 ± 0.021, 0: 0.219 ± 0.019, 5: 0.297 ± 0.019, 10: 0.350 ± 0.015, 15: 0.372 ± 0.019, 20: 0.397 ± 0.027, 25: 0.428 ± 0.024, 30: 0.453 ± 0.027). 3 independent experiments. Statistical significance was determined using 2-way ANOVA with Sidak’s multiple comparison test, *p<0.05, **p<0.01, ****p<0.001. Error bars are SEM. (C) The size of GFP-SNX17-positive puncta in 30 µm of dendritic spines at 10 and 30 min following cLTP (or mock) was quantified and normalized to the baseline for each dendrite. Mock: N=12 dendrites (10: 1.063 ± 0.079, 30: 1.087 ± 0.065); cLTP: N=12 dendrites (10: 1.132 ± 0.062, 30: 1.178 ± 0.079). 3 independent experiments. Data were analyzed using unpaired two-tailed Student’s t-test. Error bars are SEM. (D) The number of puncta containing GFP-SNX17-R-L470G in 30 µm of dendritic spines at the different time points following cLTP (or mock) was quantified and normalized to the baseline for each dendrite. Mock: N=12 dendrites (baseline: 0.206 ± 0.023, cLTP: 0.206 ± 0.022, 0: 0.208 ± 0.023, 5: 0.217 ± 0.023, 10: 0.225 ± 0.026, 15: 0.231 ± 0.028, 20: 0.242 ± 0.029, 25: 0.244 ± 0.031, 30: 0.253 ± 0.032); cLTP: N=12 dendrites (baseline: 0.231 ± 0.027, cLTP: 0.217 ± 0.024, 0: 0.228 ± 0.023, 5: 0.239 ± 0.025, 10: 0.250 ± 0.023, 15: 0.264 ± 0.025, 20: 0.283 ± 0.029, 25: 0.292 ± 0.027, 30: 0.303 ± 0.027). 3 independent experiments. Two-way ANOVA with Sidak’s multiple comparison test determined that there are no significant differences between the mock and cLTP conditions. Error bars are SEM. (E) DIV17 hippocampal neurons were treated with cLTP or HBS control (mock), washed, and incubated in HBS for 10 min following the cLTP (or mock) stimulus. Neurons were then fixed and stained for SNX17 and the Retromer marker VPS35. Arrows indicate examples of colocalization. Scale bar, 5 µm. (F) The percentage of VPS35 that colocalizes with SNX17 was quantified using Mander’s colocalization coefficient (x100). Mock-treated neurons: 55.790 ± 2.318, N=27; cLTP-treated neurons: 67.650 ± 2.688, N=30. 3 independent experiments. Data were analyzed by unpaired two-tailed Student’s t-test, **p<0.01. Error bars are SEM. (G) The percentage of Syntaxin13 that colocalizes with SNX17 was quantified using Mander’s colocalization coefficient (x100). Mock-treated neurons: 50.500 ± 1.651, N=26; cLTP-treated neurons: 59.450 ± 1.679, N=26. 3 independent experiments. Data were analyzed by unpaired two-tailed Student’s t-test, ***p<0.005. Error bars are SEM.

SNX17 recruitment to endosomal compartments requires the presence of phosphatidylinositol (3)-phosphate (PI(3)P) (Teasdale and Collins, 2011; Ghai et al., 2011), raising questions of whether cLTP-dependent changes in SNX17 localization are partially governed by alterations in PI(3)P. We transfected rat hippocampal neurons with dsRed-EEA1-FYVE, a bioprobe that potently and specifically binds PI(3)P (Singla et al., 2019). Notably, cLTP caused an increase in the number of PI(3)P puncta over time (Fig. S5A), which is similar to the cLTP-dependent increase in GFP-SNX17 puncta.

In mouse embryonic fibroblasts, approximately two thirds of the PI(3)P pool is generated by the lipid kinase VPS34 (Devereaux et al., 2013; Ikonomov et al., 2015). We previously showed that VPS34 regulates SNX17 recruitment to endosomes in HeLa cells (Giridharan et al., 2022). To determine whether PI(3)P generated by VPS34 recruits SNX17, we transiently transfected neurons with dsRed-EEA1-FYVE and GFP-SNX17, and analyzed the numbers of PI(3)P puncta and GFP-SNX17 puncta at 10 and 30 min following cLTP in the presence or absence of VPS34 inhibitor (VPS34-INH). VPS34-INH treatment caused a significant reduction in the number of PI(3)P puncta and also blocked the cLTP-dependent increase in PI(3)P puncta at 10 min. Surprisingly, 30 min after cLTP in the presence of VPS34-INH, there was a small but significant elevation in PI(3)P puncta compared to the baseline (Fig. S5B and C). This increase could potentially be due to VPS34-independent PI(3)P synthesis.

Importantly, in the absence of VPS34 inhibition, the changes in PI(3)P puncta correlate with changes in GFP-SNX17-positive puncta in dendrites (Fig. S5B and D). Moreover, SNX17 showed good colocalization with PI(3)P at the time points analyzed (baseline: 63.4 ± 3.2%, 10 min: 68.9 ± 3.3%, 30 min: 70.3 ± 3.1%) (Fig. S5E). Together, these observations suggest that PI(3)P synthesis is necessary for the increase in SNX17-positive puncta during cLTP.

### SNX17 regulates the surface levels of β1-integrin in neurons

That the SNX17 pathway is actively engaged after cLTP suggests that recycling of SNX17-dependent cargoes plays a role in the functional and/or structural plasticity necessary for LTP. In HeLa cells, many adhesion molecules, signaling receptors and solute transporters recycle via the SNX17-Retriever pathway (McNally et al., 2017). However, in neurons, most cargoes which traffic via SNX17-Retriever, have not yet been identified. We chose to test β1-integrin, which is a well-characterized SNX17-Retriever cargo in non-neuronal cells and plays several roles in regulating neuronal function and synaptic plasticity. β1-integrin plays key roles in neurite outgrowth, axon guidance, and in the formation and maintenance of synapses (Cheah and Andrews, 2018; Lilja and Ivaska, 2018; Park and Goda, 2016).

To determine if SNX17 is required for normal surface levels of β1-integrin in neurons, we transduced neurons with lentivirus carrying an shRNA targeting SNX17 and performed surface biotinylation assays. SNX17 knockdown caused a 42.4 ± 3.1 % reduction in surface β1-integrin levels (Fig. 8A and B), which suggests that β1-integrin is a SNX17 cargo in neurons. Similarly, monitoring surface-exposed β1-integrin by labeling with an antibody specific for an extracellular epitope, showed that SNX17 knockdown induced a significant loss of surface β1-integrin in neurons under basal conditions relative to control (scrambled) shRNA (Fig. 8C-D).

**Figure 8.**
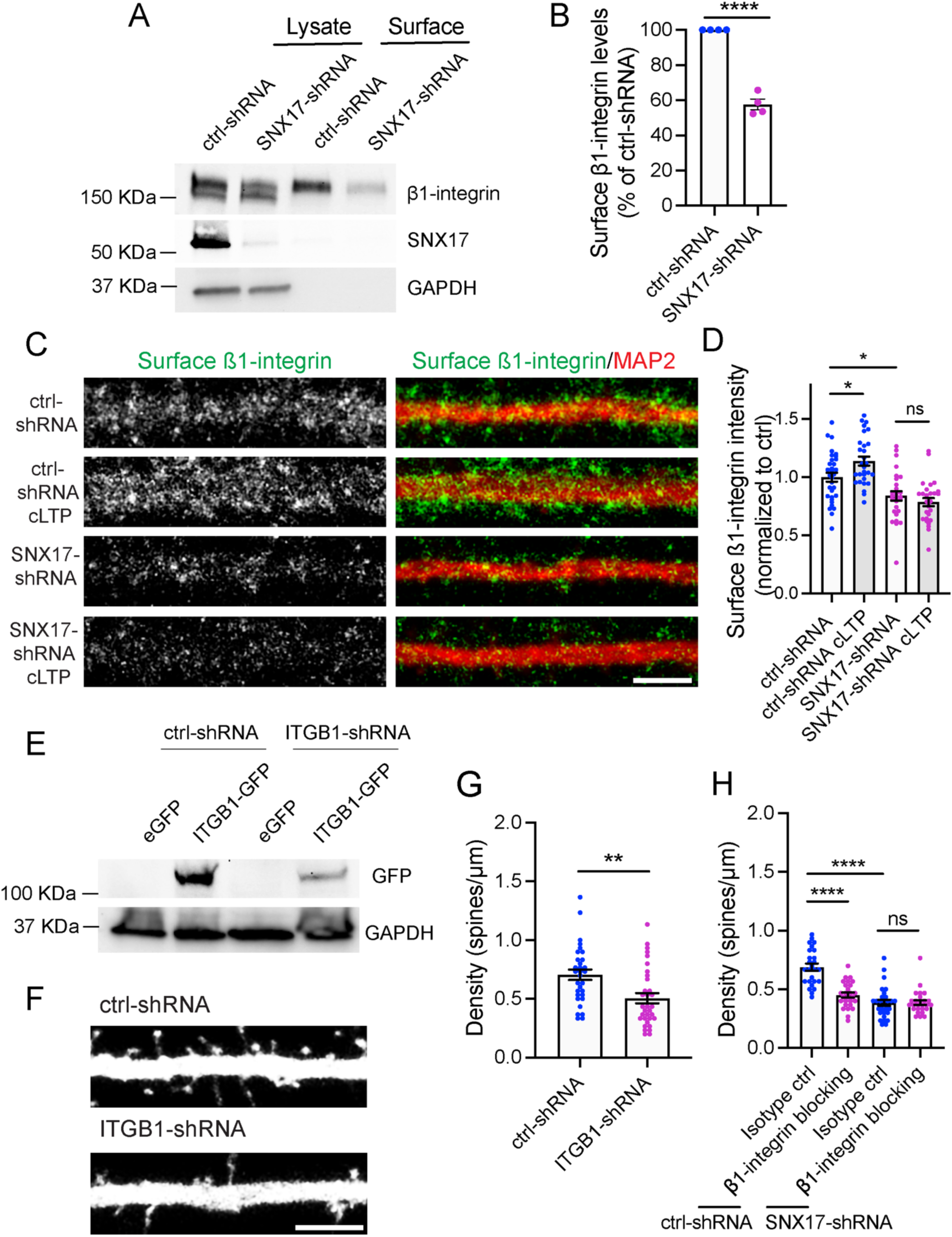
β1-integrin is a SNX17 cargo in neurons and plays a role in dendritic spine density. (A) DIV11 rat cortical neurons were infected with lentiviruses carrying scramble or SNX17 shRNAs, and the surface levels of β1-integrin were determined at DIV17 using a surface biotinylation assay. SNX17 knockdown was validated by western blotting of the lysate, and GAPDH was used as a loading control. (B) The levels of surface β1-integrin protein were quantified and normalized to total β1-integrin levels (lysate). Data are expressed as percentage of ctrl-shRNA (ctrl-shRNA: 100%, SNX17-shRNA: 57.630 ± 3.058%). N=4 independent experiments. Statistical significance was determined using unpaired two-tailed Student’s t-test, ****p<0.001. Error bars are SEM. (C) Representative confocal images of surface β1-integrin levels of DIV17 hippocampal neurons that were infected at DIV11 with lentiviruses carrying either ctrl-shRNA or SNX17-shRNA. Neurons were treated in the presence or absence of cLTP and live labeled with an anti-surface ß1-integrin antibody for 15 min, followed by fixation and immunostaining for MAP2. Scale bar, 5 µm. (D) The intensity of ß1-integrin in the first 50 µm of secondary dendrites was quantified, and values were normalized to crtl-shRNA. ctrl-shRNA: 1.000 ± 0.038, N=32 neurons; ctrl-shRNA cLTP: 1.139 ± 0.039, N=29 neurons; SNX17-shRNA: 0.839 ± 0.040, N=28 neurons; SNX17-shRNA cLTP: 0.786 ± 0.035, N=28 neurons. 3 independent experiments. Data were analyzed by one-way ANOVA with Tukey’s post hoc test, *p<0.05. Error bars are SEM. (E) Validation of a shRNA clone (V2LMM_39157, Horizon Discovery) to knock-down rat ITGB1. pGIPZ scrambled non-target (RHS4346, Horizon Discovery) was used as a control. HEK293 cells stably expressing the tet repressor (TR-HEK293) were either transfected with control-shRNA or ITGB1-shRNA in the absence or presence of eGFP or ITGB1-GFP, as indicated. 5-days post-infection, cells were treated with 1 μg/ml of doxycycline to promote the expression of eGFP or ITGB1-GFP. 24 hours later, extracts were generated and analyzed by western blot. (F) Representative confocal images of dendritic spines in DIV16 hippocampal neurons transfected at DIV12 with eGFP (filler) and either ctrl-shRNA or ITGB1-shRNA. Scale bar, 5 µm. treated with either β1-integrin blocking or iso type control antibodies 24 hours before fixation. Scale bar, 5 µm. (G) The numbers of dendritic spines in the first 30 μm of secondary dendrites were quantified. ctrl-shRNA: 0.705 ± 0.044, N=31 neurons; ITGB1-shRNA: 0.505 ± 0.043, N=33 neurons. Statistical significance was determined using unpaired two-tailed Student’s t-test, **p<0.01. Error bars are SEM. (H) Hippocampal neurons were transfected at DIV12 with eGFP (filler) and either ctrl-shRNA or SNX17-shRNA. Neurons were treated with either β1-integrin blocking or isotype control antibodies 24 hours before fixation at DIV16. The number of dendritic spines in the first 30 μm of secondary dendrites was quantified. ctrl-shRNA + isotype ctrl: 0.689 ± 0.030, N=26 neurons; ctrl-shRNA +β1-integrin blocking: 0.451 ± 0.021, N=27 neurons; SNX17-shRNA + isotype ctrl: 0.385 ± 0.026, N=28 neurons; SNX17-shRNA +β1-integrin blocking: 0.387 ± 0.020, N=26 neurons,. Statistical significance was determined using one-way ANOVA with Tukey’s post hoc test, ****p<0.001. Error bars are SEM

That SNX17 is engaged by synaptic activity, led us to test whether surface β1-integrin levels are regulated by synaptic activity in a SNX17-dependent manner. We induced cLTP and found a significant increase in the surface levels of β1-integrin. Importantly, surface β1-integrin was not altered by cLTP stimulation following SNX17 knockdown (Fig. 8C and D). These findings suggest that SNX17-Retriever-dependent recycling dynamically controls β1-integrin surface expression in hippocampal neurons.

### Blocking β1-integrin function mimics most effects of SNX17 knockdown on synaptic function

Our results suggest that the role of SNX17 in synaptic function and plasticity may be mediated in part by β1-integrin. To more directly test a role for β1-integrin in synaptic function, we used an ITGB1-shRNA construct to decrease β1-integrin levels. The ability of the shRNA to knockdown ITGB1 was first tested in HEK293 cells by western blot analysis (Fig. 8E).

We then transfected the ITGB1-shRNA plasmid into hippocampal neurons and found that 4 days after transfection, there was a 28.2% decrease in the number of dendritic spines as compared to cells transfected with scrambled control shRNA (ctrl-shRNA: 0.705 ± 0.044, ITGB1-shRNA: 0.505 ± 0.043) (Fig. 8F and G). As an orthogonal approach, we treated neurons for 24 hours with an antibody that blocks β1-integrin function (Kramár et al., 2006; Wang et al., 2018b) or an isotype control antibody, and quantified dendritic spine density. Similar to shRNA-mediated knockdown, treatment with β1-integrin blocking antibodies caused a reduction in spine density as compared to the isotype control (Fig. 8H). Moreover, the reduction in spine density following a 24 h treatment with β1-integrin blocking antibodies was similar to that observed upon SNX17 knockdown. Importantly, addition of β1-integrin blocking antibody had no further effect on dendritic spine density in neurons subjected to SNX17 knockdown (Fig. 8H), which suggests that both SNX17 and β1-integrin act in the same pathway for synapse maintenance.

Finally, we tested whether blocking β1-integrin impairs enduring functional changes in synaptic efficacy that accompany LTP. Neurons were pre-treated with blocking or control antibodies for 30 minutes, and then mock-treated or treated with cLTP stimulation. mEPSCs were recorded in the presence of the antibodies (Fig. 9A). Interestingly, the initial cLTP-dependent enhancement of synaptic function was not altered by β1-integrin blocking antibodies during the first 30 min following cLTP. However, the maintenance of cLTP-dependent changes from 30-60 min post cLTP was completely disrupted (Fig. 9B-D). This is in contrast to SNX17 knockdown, which blocked the initiation of cLTP. These findings suggest that rather than β1-integrin, other cargoes of SNX17 are required during the initiation of LTP. Together, these results reveal a role for β1-integrin in the maintenance of LTP and indicate that the roles of the SNX17-Retriever pathway in cLTP are mediated in part by β1-integrin.

**Figure 9.**
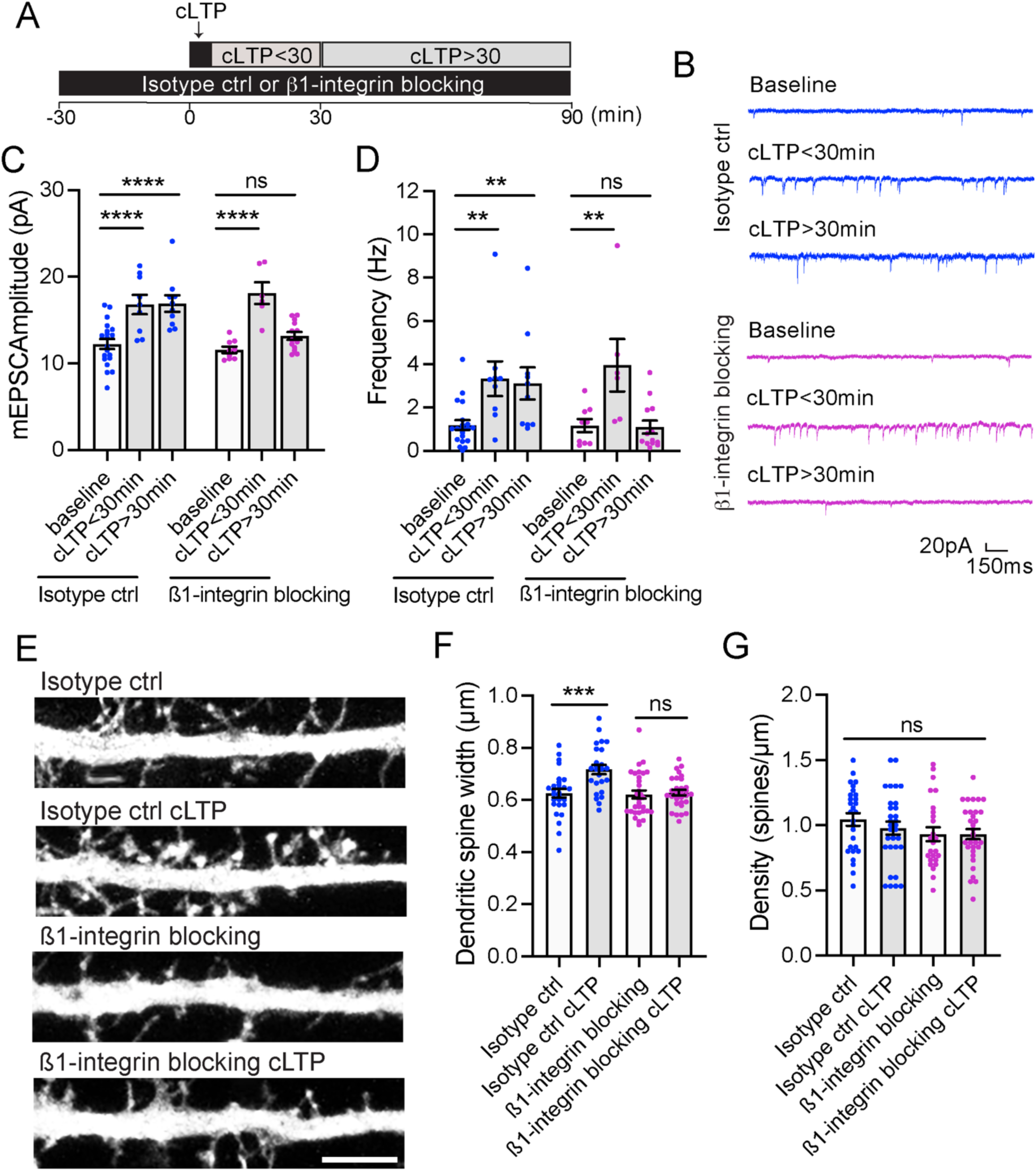
β1-integrin has roles in structural and functional plasticity during cLTP. (A) Diagram of experiment. DIV16-18 hippocampal neurons were treated with either β1-integrin blocking or isotype control antibodies for 30 min, followed by a 5 min cLTP stimulus. mEPSCs were recorded during the first 30 min after cLTP (cLTP<30) or from 30 to 90 min after cLTP (cLTP>30). (B) Examples of mEPSC recordings of neurons that were treated with isotype ctrl or β1-integrin blocking antibodies. Recordings were performed in the absence of cLTP (baseline), during the first 30 min after cLTP or from 30 to 90 min after cLTP. (C) Quantification of mEPSC amplitude. Isotype ctrl baseline: 12.240 ± 0.570, N=19 neurons; isotype ctrl cLTP<30min: 16.790 ± 1.106, N=9 neurons; isotype ctrl cLTP>30min: 16.900 ± 0.954, N=10 neurons; β1-integrin blocking baseline: 11.570 ± 0.373, N=9 neurons; β1-integrin blocking cLTP<30 min: 18.110 ± 1.247, N=6 neurons; β1-integrin blocking neurons cLTP>30 min: 13.170 ± 0.458, N=13 neurons. 3 independent experiments. Data were analyzed by one-way ANOVA with Tukey’s post hoc test, ****p<0.001. Error bars are SEM. (D) Quantification of mEPSC frequency. Isotype ctrl baseline: 1.191 ± 0.227, N=19 neurons; isotype ctrl cLTP<30min: 3.333 ± 0.800, N=9 neurons; isotype ctrl cLTP>30min: 3.110 ± 0.740, N=10 neurons; β1-integrin blocking baseline: 1.167 ± 0.296, N=9 neurons; β1-integrin blocking cLTP<30 min: 3.952 ± 1.214, N=6 neurons; β1-integrin blocking neurons cLTP>30 min: 1.096 ± 0.302, N=13 neurons. 3 independent experiments. Data were analyzed by one-way ANOVA with Tukey’s post hoc test, **p<0.01. Error bars are SEM. (E) Representative confocal images of dendritic spines in DIV16 hippocampal neurons transfected with eGFP (filler) at DIV12. Neurons were treated with ß1-integrin blocking or isotype control antibodies for 30 min, followed by a 5-min cLTP stimulus in the presence of antibodies where indicated. Neurons were further incubated in the presence of antibodies for 50 min before fixation. Scale bar, 5 µm. (F) The maximum width for each spine was quantified, and the average size of the dendritic spines in the first 30 μm of secondary dendrites was calculated. Isotype ctrl: 0.626 ± 0.016, N=28 neurons; isotype ctrl with cLTP: 0.717 ± 0.018, N=25 neurons; β1-integrin blocking: 0.621 ± 0.015, N=31 neurons; β1-integrin blocking with cLTP: 0.628 ± 0.010, N=32 neurons. 3 independent experiments. Data were analyzed by one-way ANOVA with Tukey’s post hoc test, ***p<0.005. Error bars are SEM. (G)Quantification of dendritic spine density (spines/μm). Isotype ctrl: 1.044 ± 0.048, N=28 neurons; isotype ctrl with cLTP: 0.977 ± 0.050, N=25 neurons; β1-integrin blocking: 0.931 ± 0.054, N=31 neurons; β1-integrin blocking with cLTP: 0.931 ± 0.039, N=32 neurons. 3 independent experiments. Data were analyzed by one-way ANOVA with Tukey’s post hoc test. Error bars are SEM.

Since β1-integrin regulates dynamic changes in the actin cytoskeleton (Delon and Brown, 2007; Michael and Parsons, 2020), we tested whether β1-integrin is required for structural changes in spine size during cLTP. We treated neurons with β1-integrin blocking or isotype control antibodies for 30 minutes before cLTP, added the cLTP stimulus, and fixed the neurons 50 minutes after cLTP. β1-integrin blocking or control antibodies were maintained in all the steps. Note that in contrast to the 24 h treatment with β1-integrin blocking antibodies, this short 30 min pre-treatment did not alter spine density (Fig. 9G). However, treatment with β1-integrin blocking antibodies prevented the cLTP-dependent increase in dendritic spine width (Fig. 9E and F). Taken together, these studies demonstrate that SNX17-dependent recycling is critical to maintain dendritic spine density and for the structural changes in dendritic spines that are associated with cLTP.

## Discussion

The studies reported here reveal that the SNX17-Retriever recycling pathway is localized to excitatory synapses and is actively engaged during long-lasting forms of synaptic plasticity. We found that synaptic activity engages SNX17-dependent recycling through recruitment of SNX17 and the Retriever complex to synaptic sites, in processes driven by NMDA receptor activation, CaMKII-dependent signaling, and endosomal PI(3)P. Furthermore, analysis of β1-integrin strongly suggests that the increased recruitment of SNX17 to postsynaptic sites upon cLTP results in an increase in activity-dependent recycling of SNX17 cargoes to the plasma membrane (Figure 10). Together, these studies demonstrate that SNX17-dependent recycling is critical to maintain excitatory synapses and for the structural changes in dendritic spines that are associated with cLTP. Future studies will be required to further test these findings *in vivo*.

**Figure 10.**
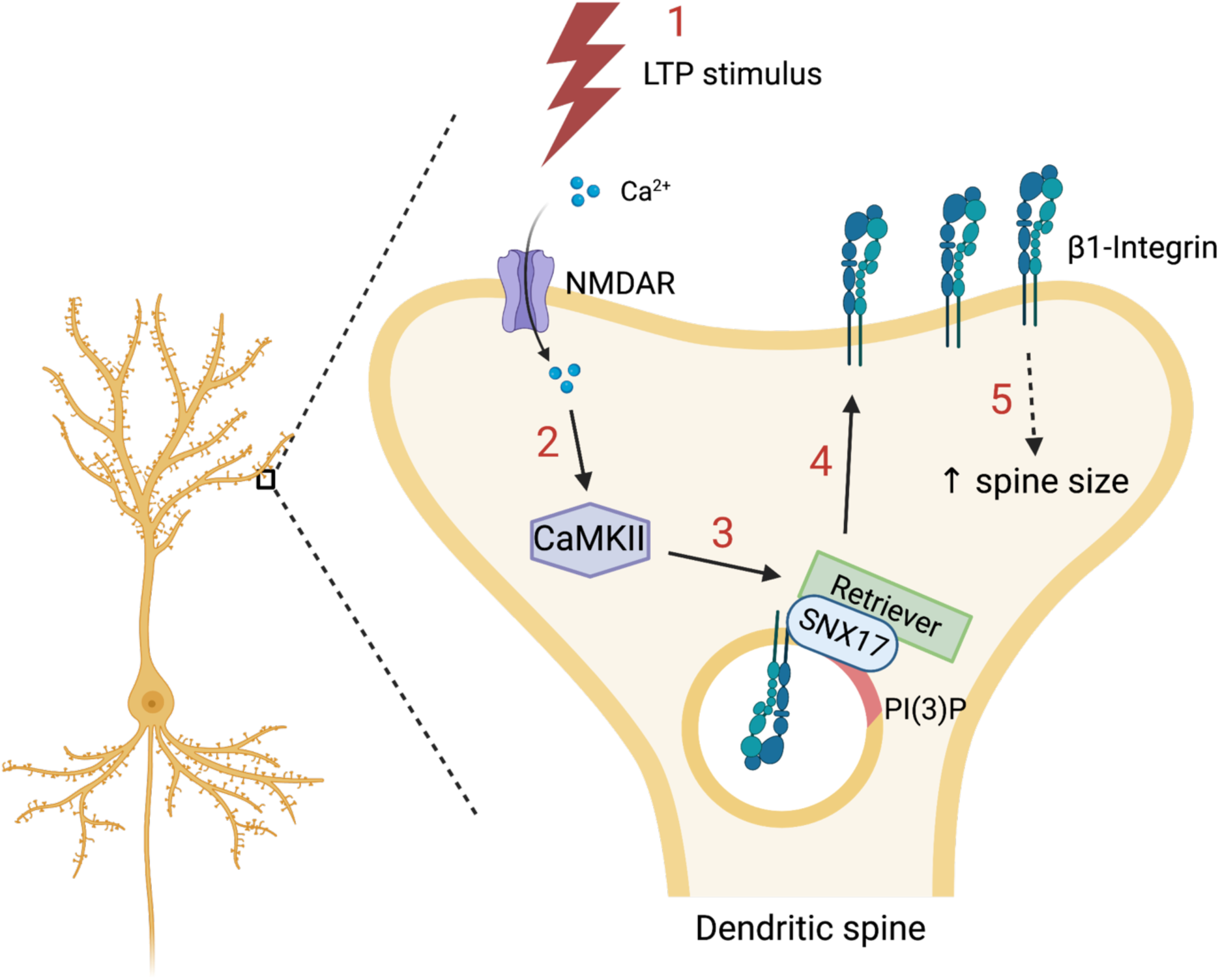
Model of SNX17-mediated modulation of synaptic structure and function. The SNX17-Retriever pathway is required for dendritic spine maintenance and for the cLTP-dependent increase in dendritic spine size. Glycine-mediated cLTP (1) stimulates calcium entry through the NMDA receptor, which activates the CaMKII pathway (2). CaMKII activation is necessary and sufficient to promote the recruitment of SNX17 and the Retriever complex to dendritic spines (3), and activates the recycling of β1-integrin from endosomes to the plasma membrane (4). The surface levels of β1-integrin increase during cLTP and promote dendritic spine growth (5). Endosomal PI(3)P increases upon cLTP and may help with the recruitment of SNX17 to synapses. Created with BioRender.com.

Importantly, these key roles for SNX17 at synapses may contribute to the neurological defects associated with mutations in subunits of this recycling pathway. Indeed, several mutations associated with Ritscher-Schinzel syndrome cause a decrease in the levels of Retriever subunits (Otsuji et al., 2022; Kato et al., 2020) or CCC subunits (Starokadomskyy et al., 2013) which interferes with the normal function of these complexes.

In neurons, multiple types of proteins are dynamically exposed at the cell surface, and undergo rapid changes in response to synaptic activity (van Oostrum et al., 2020). Among the best characterized is the LTP-dependent recycling of AMPARs from intracellular compartments, which is dependent on the SNX27 pathway (Temkin et al., 2017; Loo et al., 2014; Hussain et al., 2014; Wang et al., 2013). Our data reveal that SNX17-dependent recycling is also engaged during LTP, which likely also plays a key role in activity-dependent changes in the surface proteome.

We find that SNX17 localization is dynamically regulated by synaptic activity in response to NMDA receptor activation and CaMKII signaling. These findings establish a previously unknown link between the SNX17 pathway and neuronal-specific signaling. How increased synaptic activity triggers SNX17 recruitment to endosomes remains unclear. A variety of scenarios are possible. LTP activates signaling pathways that involve phosphorylation and dephosphorylation events (Jing et al., 2016). Notably, the ability of several SNX proteins to bind membranes in non-neuronal cells is regulated by phosphorylation (Lenoir et al., 2018; Mao et al., 2021), and this is likely to occur in neurons as well. While further work will be required to address this possibility, it is tempting to speculate that LTP-mediated posttranslational modifications may promote SNX17 recruitment to endosomes.

In addition, SNX17 binding to membranes is regulated by the simultaneous interaction with the phosphoinositide PI(3)P through its PX domain and with cargo proteins via its FERM domain (Ghai et al., 2013). PI(3)P depletion reduces neurotransmission by interfering with synaptic vesicle endocytosis (Liu et al., 2022). Here, we find that PI(3)P levels increase upon cLTP induction, and blocking this increase by inhibition of VPS34 decreases the cLTP-dependent recruitment of SNX17 to endosomal compartments. In the presence of VPS34-INH, there was a partial increase in PI(3)P levels 30 min after the cLTP stimulus, which may be due to the presence of additional sources of PI(3)P. Note that while VPS34 is responsible for a major portion of the PI(3)P pool (Devereaux et al., 2013; Ikonomov et al., 2015), class II phosphatidylinositol 3-kinases and INPP4 phosphatases also contribute to PI(3)P levels (Heng and Maffucci, 2022; Gozzelino et al., 2020; Burke et al., 2022). The sources of PI(3)P at synapses are currently unknown, but our finding that PI(3)P levels increase 30 min after cLTP even in the presence of VPS34-INH suggests that additional sources of PI(3)P are induced by cLTP. Importantly, phosphoinositide levels have been shown to change dynamically during some forms of synaptic plasticity (McCartney et al., 2014). In addition to PI(3)P, it is likely that the appearance of other interactors at specific membrane compartments may help to recruit SNX17 to these compartments during cLTP.

We find that, similar to SNX17, the Retriever complex is dynamically recruited to dendritic spines upon cLTP and that the interaction of SNX17 with Retriever is necessary for the cLTP-dependent recruitment of SNX17 to synapses, as well as for the increase in SNX17-positive puncta. Dynamic changes in phosphoinositide levels may also be involved in the recruitment of Retriever subunits (Giridharan et al., 2022).

Our studies also reveal that the roles of SNX17 in synaptic function are, in part, mediated by its cargo β1-integrin. These findings are consistent with previous studies reporting a critical role for β1-integrin in neuronal function and plasticity (Cheah and Andrews, 2018; Lilja and Ivaska, 2018; Park and Goda, 2016; Orr et al., 2022). We found that SNX17 knockdown results in lowered levels of surface exposed β1-integrin and prevents cLTP. However, while reducing SNX17 levels prevents the initiation of the cLTP response, interfering with β1-integrin function blocks the maintenance but not the initial synaptic enhancement following cLTP induction. This finding suggests that, in addition to regulating surface levels of β1-integrin, SNX17 likely regulates synaptic function via control of additional cargoes that recycle from endosomes to the cell surface. For example, LRP1, another SNX17 cargo, is critical for brain lipid metabolism and neuronal integrity (Liu et al., 2010). SNX17 also regulates ApoER2 recycling and Reelin signaling (Sotelo et al., 2014), which cooperate to enhance LTP in the mouse hippocampus (Weeber et al., 2002). There are likely to be other cargoes that rely on SNX17 that are essential for the postsynaptic changes that are induced during cLTP. In this context, gaining a full understanding of the SNX17 proteome in neurons will be critical to fully understand the roles of SNX17 in neurons.

Altered endosomal recycling has been implicated in neurodegenerative diseases, including Alzheimer’s (Small et al., 2005, 2017) and Parkinson’s disease (Vilariño-Güell et al., 2011; Zimprich et al., 2011; Zavodszky et al., 2014; McGough et al., 2014). The critical roles of SNX17 in neurons open the possibility that it may contribute to altered receptor recycling in neurodegeneration. Importantly, upregulating the SNX27-Retromer pathway has shown therapeutic potential for Alzheimer’s disease (Li et al., 2020). Similarly, the SNX17-Retriever pathway may also provide a novel therapeutic target for Alzheimer’s disease and related neurodegenerative disorders.

## Material and methods

### DNA constructs and site-directed mutagenesis

To knock-down SNX17, we used the MISSION shRNA lentiviral plasmids pLKO.1-puro with shRNA target sequence CCAGATGACTTGATCGGATAT (TRCN0000190340, Millipore Sigma) and pLKO.005-puro with shRNA target sequence GTTGGCCTGAACCTGCTTTAT (TRCN0000382281, Millipore Sigma), as clones 1 and 2, respectively. To knock-down VPS26C, a pGIPZ vector with shRNA target sequence TAATCTTGATGTCCAGAGT (V3LMM_455807, Horizon Discovery) was utilized. To knock-down ITGB1, a pGIPZ vector with shRNA target sequence TTCTTTATAGTTTGAGAGC (V2LMM_39157, Horizon Discovery) was utilized. MISSION pLKO.1 scrambled non-target shRNA SHC002 (Millipore Sigma) was used as a control for SNX17 knockdown constructs, and pGIPZ scrambled non-target (RHS4346, Horizon Discovery) was the control for the shRNAs to VPS26C and ITGB1.

pmCherry-C1 (cat. No. 632524) and pAcGFP-N1 (cat. No. 632469) were purchased from Clontech. dsRed-EEA1-FYVE was a generous gift from Dr. Daniel D. Billadeau (Mayo Clinic) and has been previously described in (Singla et al., 2019). pRK5-HA-CaMKII-T286D was a generous gift from Dr. Gentry Patrick (University of California San Diego).

pCDNA4:TO was purchased from Invitrogen, and was used to generate the pCDNA4:TO-GFP_NoStop vector (without a stop codon after GFP). To build this vector, GFP was amplified from pAcGFP-N1 with oligos 5*′*- AAAGGTACCATGGTGAGCAAGGGCGAGGAG-3*′* and 5*′*- AAGGGCCCTTACTTGTACAGCTCGTCC-3*′*. The resulting GFP construct was digested with KpnI and ApaI restriction enzymes (New England Biolabs), and ligated into an ApaI- and KpnI-digested pCDNA4:TO vector (Invitrogen) using the T4 ligase (New England Biolabs).

To generate GFP-SNX17, SNX17 was amplified from rat whole brain cDNA with oligos 5*′*- CAAAATGGCGAACTGGGCTG-3*′* and 5*′*-TCTCCTCTTGGGTAGAGGGC-3*′*, and a DNA fragment with a 5*′* KpnI site containing GFP and an 18 base pair GS linker was ordered from Twist Bioscience. To add regions of overlap, SNX17 was amplified with primers 5*′*- CAGGGGGTGGAAGCGGTGGTCACTTTTCCATTCCTGAAACC-3*′*and 5*′*- GCTGATCAGCGGGTTTAAACGGGCCCTTACAGATCCTC-3*′*. The pCDNA4:TO vector was digested with ApaI and KpnI, and NEBuilder HiFi DNA Assembly Cloning Kit (New England Biolabs) was used to assemble the digested vector with the GFP and SNX17 constructs. This generated the vector pTO-GFP-SNX17.

To create a low expression construct for live-cell imaging, GFP-SNX17 was PCR-amplified from pTO-GFP-SNX17 with primers 5’-AAAGGTACCATGGTGAGCAAGGGCGAGG-3’ and 5’-AAAGAATTCTTACAGATCCTCATCTCC-3’, digested with KpnI and EcoRI restriction enzymes, and subcloned into a modified pAAV vector. To generate this vector, pAAV-hSyn1-mNeonGreen (Addgene plasmid 99135) was digested with XbaI and KpnI to release the hSyn1 promoter. The resulting vector was ligated to the human neuron-specific enolase promoter (hENO2), which was extracted from Addgene plasmid 11606 by PCR amplification to add XbaI and KpnI flanking sites with oligos 5’- AAATCTAGATATGCAGCTGGACCTAGGAGAGAAGCAG-3’ and 5’- AAAGGTACCCGGTGGTAGTGGCGGTGGCGGTGGCGGTGG-3’. This resulted in the pAAV-eno-mNeonGreen vector. mNeonGreen was then released by digestion with KpnI and EcoRI, and the purified plasmid was used for the ligation reaction. This generated the pAAV- eno-GFP-SNX17.

pTO-GFP-SNX17 was used to generate an shRNA resistant version of GFP-SNX17 by introducing three silent mutations into the target sequence of the SNX17 shRNA clone 1 construct using the Q5-site directed mutagenesis kit (New England Biolabs). Specifically, the original sequence 5’-CCAGATGACTTGATCGGATAT-3’ was mutated to 5’- CCTGACGACTTAATCGGATAT-3’ with oligos 5’- ACTTATGATAGACGGTTTTTCGC -3’ and 5’- CGTCACCACTACGTGAACCATCAC-3’. This generated the pCDNA4:TO -GFP- SNX17-R vector.

pCDNA4:TO -GFP-SNX17-R was used to generate the L470G mutant by site-directed mutagenesis with oligos 5’-AGATGAGGATGGGTAAGGCCCGTTTAAACC-3’ and 5’- CCAATGCCCTCGAAGGCG-3’. To generate a low expression construct for live-cell imaging, the resulting construct was PCR amplified with primers 5’- CCGCCACTACCACCGGGTACCACCATGTGAGCAAGGGC-3’ and 5’- TATCGATAAGCTTGATATCGAATTCTTACCCATCCTCATCTCCAATG-3’, and subcloned into the KpnI- and EcoRI-digested pAAV-eno-mNeonGreen vector.

To generate VPS26C-GFP, VPS26C was amplified from rat whole brain cDNA with oligos 5*′*- GCCTTTGTGGATAATCCGAGATG-3*′* and 5*′*-CCACTCTGTCCCATTCCTGC-3*′*. A synthetic DNA fragment containing an 18 base pair GS linker and GFP between the restriction sites for ApaI (5*′*) and KpnI (3*′*) was ordered from Twist Bioscience. Both the synthetic DNA fragment and the pCDNA4:TO-GFP vector were digested with ApaI and KpnI, and the resulting constructs were ligated using the T4 ligase to generate pCDNA4:TO-GFP-linker. This vector was linearized by digestion with KpnI. Regions of overlap to pCDNA4:TO-GFP-linker were added to VPS26C by PCR amplification with primers 5*′*- TTTAAACTTAAGCTTGGTACCATGGGGACTACTCTGGAC -3*′*and 5*′*- CCACCCCCTGAACCGCCCCCCGTCCGACAGAGCTTCAG -3*′*. The resulting construct was ligated to pCDNA4:TO-GFP-linker using NEBuilder HiFi DNA Assembly.

To generate ITGB1 with an internal GFP tag (Huet-Calderwood et al., 2017), ITGB1 was amplified from rat whole brain cDNA with oligos 5*′*-GAGACCATCCGAGAAGCCG-3*′* and 5*′*- AGAGCCCCAAAGCTACCCTA-3*′*. Two fragments containing nucleotides 1-303 and 304-2,397 were PCR-amplified from ITGB1 with oligos 5*′*- TTTAAACTTAAGCTTGGTACACCATGAATTTGCAACTGGTTTTC-3*′*and 5*′*- CGCCAAACTCCCCTTTGCTGCGATTGGTG-3*′* for fragment 1, and 5*′*- CGGACTGGAAATGGCAGAGAAGCTCCGG-3*′* and 5*′*- TCAGCGGGTTTAAACGGGCCTCATTTTCCCTCATACTTCGGATTG-3*′* for fragment 2. GFP was amplified from pCDNA4:TO-GFP_NoStop using primers that added 5*′* and 3*′* 12-nucleotide GS linkers: 5*′*-GAGTTTGGCGGTATGGTGAGCAAGGGCGAGGAGC-3*′* and 5*′*- TTCCAGTCCGCCCTTGTACAGCTCGTCCATGCCG-3*′*. The resulting fragment was PCR-amplified with primers 5*′*-CAGCAAAGGGGAGTTTGGCGGTATGGTG-3*′* and 5*′*- TCTCTGCCATTTCCAGTCCGCCCTTGTAC-3*′* to add regions of overlap. The pCDNA4:TO vector was digested with ApaI and KpnI, and NEBuilder HiFi DNA Assembly was used to assemble the digested vector with the 2 ITGB1 fragments and GFP.

DNA was prepared from bacterial cultures grown at 37°C using a Midiprep kit (PureYield Plasmid Midiprep System, Promega) according to the manufacturer’s instructions. In all cases, the identity of the constructs was verified by sequencing the entire coding region.

### Cell culture and transfection

Timed pregnant Sprague-Dawley rats were obtained from Charles River Laboratories (Wilmington, MA). All experimental protocols were approved by the University of Michigan Committee on the Use and Care of Animals. Dissociated hippocampal neuron cultures, prepared from postnatal day 1–2 rat pups of either sex, were plated at a density of 50-70K cells in poly-D-lysine-coated 35-mm glass bottom Petri dishes (Mattek), as previously described (Henry et al., 2017). Neurons were maintained for the indicated days in vitro (DIV) at 37 °C and 5% CO_2_ in growth medium [Neurobasal A (Invitrogen) supplemented with B27 (Invitrogen) and Glutamax (Invitrogen)].

For western blot experiments, rat cortical neurons were prepared from postnatal day 1–2 rat pups of either sex and plated on 35-mm dishes at a density of 6,000,000 cells per dish.

Rat2 and Hek293 cells were cultured in 100-mm dishes in full medium (DMEM containing 10% fetal bovine serum and high glucose) at 37°C in 5% CO2. Cells were tested for mycoplasma using LookOut Mycoplasma PCR Detection Kit (Sigma Aldrich).

Neurons were transfected at DIV12-14 using Lipofectamine 2000 (Invitrogen) according to the manufacturer’s recommendations and used 4-5 days after shRNA transfection or 1 day after transfection of other constructs.

### Lentivirus shRNA knockdown of rat SNX17

To test shRNA clones in Rat2 cells, lentiviruses for SNX17 clones 1 and 2, VPS26C, pLKO.1 scrambled and pGIPZ scrambled were generated. To produce lentiviral particles, HEK293T cells were transfected with packaging vectors pMD2.G and psPAX2 along with relevant shRNA using Fugene 6 transfection reagent (Promega). Viral particles were harvested after 48 hours in DEMEM/ 40% FBS, aliquoted and stored at -80°C. For infection, Rat2 cells grown on two 35 mm dishes were treated with the viruses at a multiplicity of infection (MOI) of 5. After overnight incubation, cells were treated with 2 mg/ml puromycin. After two days of infection, cells from two 35 mm dishes were transferred to a 100 mm dish and maintained in puromycin containing media for another three days prior to western blot analysis.

For experiments with cultured rat neurons, transduction-ready viral particles of SNX17 shRNA clone 1 and pLKO.1 scrambled shRNA were produced by the University of Michigan Vector Core with a concentration of 10^7^ transduction units per ml. Neurons were infected at a MOI of 2, without polybrene. After overnight incubation, virus-containing medium was replaced with saved conditioned medium. Experiments were performed after 6 days of lentivirus transduction.

### Generation of the TR-HEK293 cell line

To validate shRNA-resistant GFP-SNX17, VPS26C-shRNA and ITGB1-shRNA, we used a HEK293 cell line that stably expresses the tet repressor (TR-HEK293). This cell line was generated by culturing cells in 10-cm dishes to 70% confluence, then transfecting them with 5 μg of the pCDNA6:TR vector (Invitrogen; this vector contains a blasticidin resistance cassette) using Lipofectamine 2000, according to the manufacturer’s instructions. 24 hours after transfection, the medium was replaced with fresh medium containing 5 μg/ml blasticidin (ThermoFisher Scientific) and the cells were cultured for 10 days. Cells were then harvested and sorted into a 96-well plate (1 cell/well) at the University of Michigan Flow Cytometry Core. 6 clonal lines were expanded and transient transfection with the pTO-Egfp vector (obtained by subcloning eGFP into the pCDNA4:TO vector at the KpnI and ApaI sites), was performed. Transfected cells were tested with 1 μg/ml doxycycline (Sigma) added for 12 hours to allow for eGFP expression, and a clone was chosen that had no detectable baseline eGFP.

HEK293-TR were co-transfected with the shRNA to be validated together with a pCDNA4:TO vector expressing a GFP-fused version of the shRNA target. This allows for the detection of the protein expression levels with an anti-GFP antibody. Note that we did not find commercially available antibodies that detect endogenous rat VPS26C. While shRNA expression occurs for 7 days, doxycycline addition for the last 24 h allows for the controlled expression of the GFP-fused target.

### Chemically-induced LTP

Under baseline conditions, neurons were incubated in HEPES-buffered saline (HBS) containing (in mM) the following: 119 NaCl, 5 KCl, 2 CaCl2, 2 MgCl2, 30 Glucose, 10 HEPES, pH 7.4. Pharmacological induction of LTP in cultured hippocampal neurons was achieved via brief (5 min) exposure to a Mg^2+^ −free HBS solution supplemented with: 0.4 mM Glycine (Fisher), 0.02 mM Bicuculline (Abcam), and 0.003 mM Strychnine (Tocris). Neurons were immediately washed with warm HBS after glycine stimulation and imaged or fixed at the indicated time points.

To evaluate the effect of compounds on cLTP, the following reagents were used: DMSO, 10 μM BAPTA-AM (Calbiochem), 100 μM D-APV (Sigma), 10 μM nifedipine (EMD Millipore), 10 μM AIP (Sigma-Aldrich), 2 μM KT5720 (Millipore-Sigma), 10 μM U0126 (LC Laboratories) and 1 μM VPS34-INH (EMD Millipore). Neurons were pretreated for 30 min before cLTP, and the compounds were maintained during the whole experiment.

### Electrophysiology

Whole-cell patch-clamp recordings of mEPSCs were made with a MultiClamp 700B amplifier using cultured hippocampal neurons bathed in HEPES-buffered saline [HBS; 119 mM NaCl, 5 mM KCl, 2 mM CaCl2, 2 mM MgCl2, 30 mM glucose, 10 mM HEPES (pH 7.4)] plus 1 μM TTX and 10 μM bicuculline. The pipette internal solution contained 100 mM cesium gluconate, 0.2 mM EGTA, 5 mM MgCl2, 40 mM HEPES, 2 mM Mg-ATP, 0.3 mM Li-GTP, and 1 mM QX314 (pH 7.2), and had a resistance of 3–5 MΩ. mEPSCs were analyzed offline using MiniAnalysis (Synaptosoft) and Clampfit (Molecular Devices).

### Immunofluorescence and labeling of surface exposed β1-integrin

Primary antibodies used were SNX17 rabbit polyclonal antibody (pAb) (1:200, HPA043867, Atlas Antibodies), SNX17 mouse monoclonal antibody (mAb) (1:50, sc-166957, Santa Cruz Biotechnology), VPS35L rabbit pAb (1:200, Daniel D. Billadeau, previously described in (Phillips-Krawczak et al., 2015), COMMD1 rabbit pAb (1:200, 11938-1-AP, Proteintech), VPS35 goat pAb (1:200, ab10099, Abcam), PSD-95 mouse mAb (1:200, MAB1596, Millipore Sigma), EEA1 mouse mAb (1:200, 48453, Cell Signaling), vGLUT1 guinea pig pAb (1:2000, AB5905; Millipore Sigma), Syntaxin 12/13 rabbit pAb (1:200, 110 132, Synaptic Systems), MAP2 mouse mAb (1:200, M4403, Millipore Sigma), and β1-integrin clone HM β1-1 Armenian hamster mAb (1:50, 102202, Biolegend). Secondary antibodies were conjugated to Alexa Fluor 405, 488, 594 and 647 (1:1000, Invitrogen).

Neurons were fixed with 4% paraformaldehyde/4% sucrose in PBS with 1 mM MgCl_2_ and 0.1 mM CaCl_2_ (PBS-MC) for 15 min, permeabilized with 0.1% Triton X-100 for 10 min and blocked with 2% BSA for 1 h. Neurons were then incubated with primary antibody in blocking buffer overnight followed by washes in PBS-MC and incubation with secondary antibodies for 1 h. For staining with the PSD95 antibody, 4% paraformaldehyde/4% sucrose was replaced by 2% paraformaldehyde/2% sucrose.

To label surface β1-integrin, neurons were incubated live with 10 µg/ml of β1-integrin antibody clone HM β1-1 (102202, Biolegend) for 15 min at 37 °C, fixed with 4% paraformaldehyde and 4% sucrose for 20 min, blocked with 2% BSA in PBS-MC for 1 h, and incubated with fluorescent secondary antibody (goat anti-Armenian hamster 488). Cells were then permeabilized with 0.1% Triton X-100 for 10 min and blocked before incubation with MAP2 antibody and, later, donkey anti–mouse-647 antibody.

### Image acquisition and analysis

Images were acquired with a Leica SP5 or Leica Stellaris 5 confocal microscope under an oil immersion 63X objective (z-series, 0.4 μm intervals). Images were analyzed and processed using ImageJ.

For colocalization studies, individual z-stack images corresponding to the neurite center were acquired. The JACoP plugin of ImageJ was used for the quantification of colocalization of SNX17, VPS35L or COMMD1 with endolysosomal markers. After thresholding, the percentage of colocalization was obtained by calculating the Manders’ coefficients (M1 for the proportion of protein overlapping with endolysosomal marker), and the percentage of colocalization was obtained by M1 x 100. To analyze the colocalization of SNX17, VPS35L and COMMD1 with active synapses, a mask of the overlap between PSD95 and vGLUT1 was first generated, and the colocalization of this mask with the protein of interest was calculated using JACoP.

Dendritic spine numbers and morphology were quantified manually by an observer blind to the experimental conditions in 30 or 50 μm segments, as indicated, of dendrites after the first dendritic branchpoint. Spine width was quantified by placing the line tool (in ImageJ) over the maximum spine head width, and the number of spines was counted and normalized to the length of the dendrite.

The classification of dendritic spine types was done as previously described (Henry et al., 2017). The following parameters were measured using the line tool in ImageJ: head width, length and neck width, and used to establish 5 different classes of dendritic spines. Filopodia are defined as protrusions with a length of at least 5 μm. Mushroom spines are defined as having a head to neck ratio equal to or greater than 2.5. Flat spines are defined as having a head width to length ratio of at least 1. Thin spines are defined as having a length to neck width ratio greater than or equal to 3. A spine that does not satisfy any of the previous conditions is considered a stubby spine. The class of each spine is determined by checking against these conditions sequentially.

To analyze the surface levels of β1-integrin, the integrated density in 50 μm segments of dendrites in MAP2-stained cells was measured.

### Live imaging of GFP-SNX17 and analyses

For observation of GFP-SNX17 or GFP-SNX17-R-L470G dynamics, neurons were transfected with GFP-SNX17 (or the L470G mutant construct) and mCherry 24 hours before the experiment. Media was replaced with fresh HBS 10 minutes before imaging. Neurons were then treated with cLTP or with a mock cLTP stimulation with HBS for 5 min, followed by washing with warm HBS and addition of new HBS media.

Images were acquired with an AiryScan Zeiss LSM880 scanning confocal microscope with a 63× Plan-Apo oil immersion objective. Temperature was maintained at 37°C using a microscope incubation chamber. Z-stacks (0.5 μm intervals) were acquired before adding the cLTP stimulus (baseline), 2.5 min into the cLTP stimulus (during cLTP), directly after removing the stimulus (time 0), and every 5 min till 30 min after cLTP.

mCherry was used to identify and draw individual dendritic spines in the first 30 μm of one secondary dendrite per neuron. The mean intensity of GFP-SNX17 in individual spines that could be detected in the different time points was measured using ImageJ. The intensity at the different time points after cLTP was normalized to the baseline intensity for each spine.

The total number of GFP-SNX17-positive puncta in the first 30 μm of a secondary dendrite per neuron was quantified for each time point using ImageJ software, and data were normalized to the baseline number of puncta for each neuron.

### ß1-integrin blocking studies

To determine the effect of ß1-integrin in spine density, cells were treated with function-blocking anti-ß1-integrin monoclonal antibodies (MAB1987Z, Millipore Sigma) at a concentration of 10 µg/ml and fixed 24 hours later. Control cells were treated with 10 µg/ml of IgG2a isotype control antibodies (MABC004, Millipore Sigma). For cLTP studies, cells were pre-incubated for 30 minutes with ß1-integrin blocking or isotype control antibodies at a concentration of 10 µg/ml, followed by cLTP in the presence of antibodies. Cells were then washed, and fresh HBS media with antibodies was added. Neurons were either used for electrophysiology or fixed 50 minutes later for spine size analysis.

### Surface biotinylation assays

All solutions were pre-chilled to 4°C and all steps were carried out in ice to prevent internalization of surface β1-integrin. Cells were washed with wash buffer (PBS containing 2.5 mM MgCl_2_ and 1 mM CaCl_2_) and incubated with 0.2 mg/ml NHS-SS-Biotin (Pierce) for 15 min. Neurons were then washed with wash buffer before being quenched in quenching buffer (50 mM Triz, 100 mM NaCl, pH 7 for 10 min. Neurons were lysed in 2% Triton X-100 containing protease inhibitors (Roche). A BCA assay (Pierce) was used to determine protein concentrations, and 3 mg of protein lysate was incubated with Dynabeads MyOne Streptavidin C1 (ThermoFisher Scientific) for 1 h at 4°C. Western blot analysis was performed on 50% of the total of each immunoprecipitate and 50 µg of each lysate.

### Cell extracts and western blot analysis

Cells were collected and lysed in RIPA buffer (Pierce) containing protease inhibitors (Roche). Extracts were sonicated, boiled, and centrifuged at 10,000 g for 10 min. Protein concentrations were determined using a commercial BCA assay (Pierce), and equal amounts of protein were loaded into Mini-PROTEAN TGX Precast gels (Bio-Rad) and transferred to nitrocellulose membranes. Membranes were blocked with Tris-HCl-buffered saline containing 5% BSA and 0.1% Tween and probed with primary antibodies in blocking buffer overnight at 4°C. Primary antibodies used included SNX17 rabbit pAb (1:1000, HPA043867, Atlas Antibodies), SNX17 mouse mAb, VPS35L rabbit pAb (1:1000, Daniel D. Billadeau), COMMD1 rabbit pAb (1:1000, 11938-1-AP, Proteintech), GFP rabbit mAb (1:1000, Ab32146, Abcam), mouse/rabbit Integrin β1 goat pAb (1:1000, AF2405, R&D Systems) and GAPDH rabbit mAb (1:2000, 2118, Cell Signaling). Following washes in TBS containing 0.1% Tween-20, the blots were incubated with horseradish peroxidase conjugated secondary antibodies and developed using chemiluminescence with an Amersham ECL western blotting detection reagent according to the manufacturer’s instructions (Cytiva). Chemiluminescence signals were detected using a BioRad ChemiDoc Imaging system. Immunoblots were analyzed using ImageLab software.

### Statistical analyses

All experiments were repeated at least 3 times. All data are expressed as means ± S.E.M. Statistical tests and the size of the samples are described in the respective figure legends. Microsoft Excel software was used for calculations, and the results were plotted and analyzed using GraphPad Prism 9.

## Acknowledgements

We thank Cindy Carruthers and Christian Althaus for preparing neuronal cultures. We thank the members of the Weisman and Sutton labs for their insights and suggestions. We thank Carole Parent for giving us access to her AiryScan Zeiss LSM880 scanning confocal microscope. We also thank Daniel D. Billadeau for providing antibodies for VPS35L and the dsRed-EEA1-FYVE construct. We thank Rosalyn Adam and Viviana Gradinaru for the Addgene plasmids used in this study, as well as Gentry Patrick for providing the pRK5-HA-CaMKII-T286D construct. This work was supported by R01-NS129198 to L.S.W. and M.A.S., R01-NS099340 and R01-NS064015 to L.S.W., R01-NS097498 to M.A.S, and R21-NS125449, R21-MH127485 and R01-NS116008 to S.I., and by the University of Michigan Protein Folding Diseases Fast Forward Initiative. P.R.R. was in part supported by a Michigan Life Sciences Postdoctoral Fellowship (University of Michigan), by a NIH/NIA Michigan Alzheimer’s Disease Research Center grant P30AG072931 and the University of Michigan Alzheimer’s Disease Center (Berger Endowment). T.T. was supported by a Parents and Researchers Interested in Smith-Magenis Syndrome (PRISMS) post-doctoral fellowship. A.C. was supported by a National Institutes of Health (NIH) National Research Service Award (NRSA) fellowship (18-PAF03228).

