## Supplementary figures for "Recruitment of the SNX17-Retriever recycling pathway regulates synaptic function and plasticity"

**Figure S1**

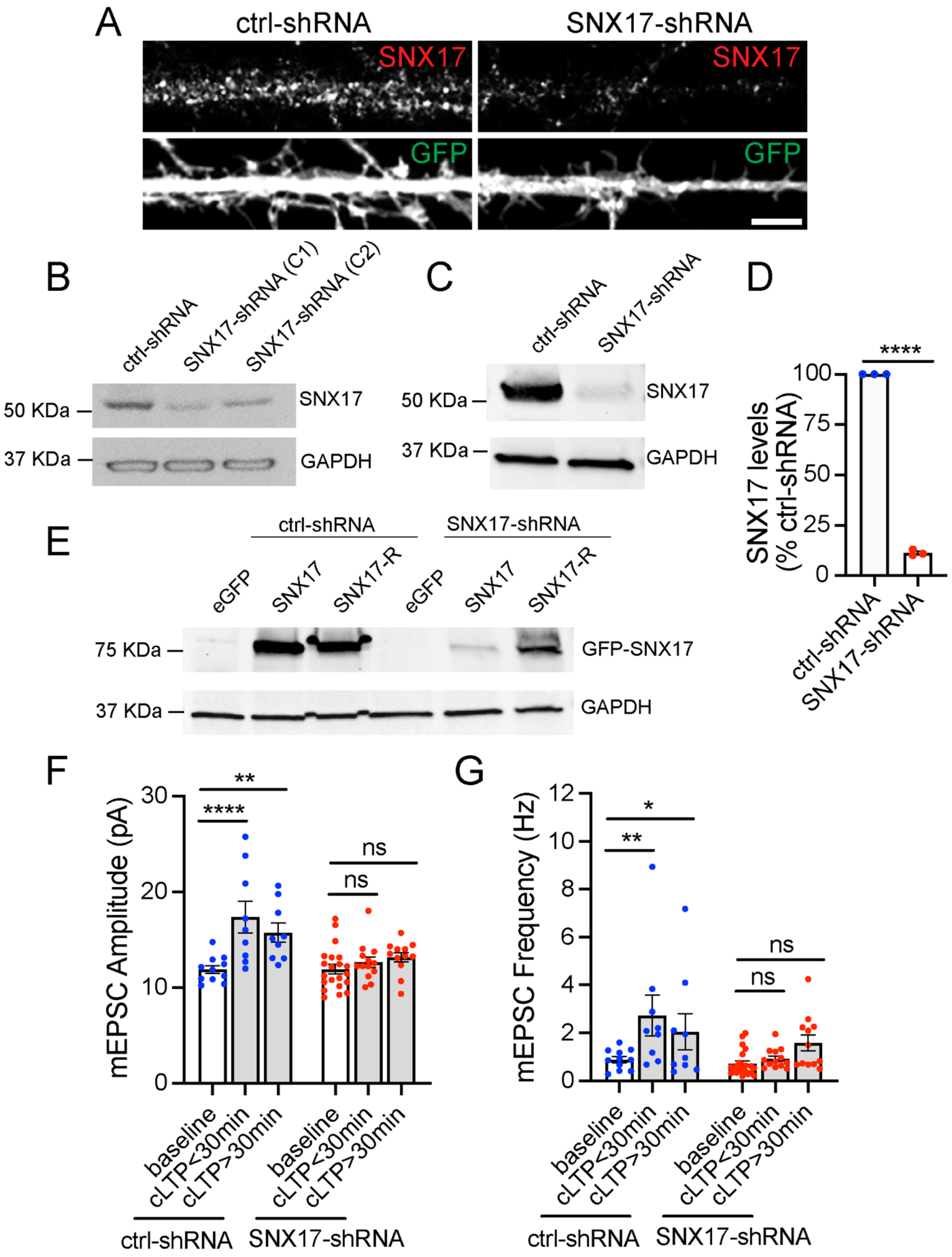

A
ctrl-shRNA
SNX17-shRNA
SNX17
GFP
B
ctrl-shRNA
SNX17-shRNA (C1)
SNX17-shRNA (C2)
50 KDa
37 KDa
SNX17
GAPDH
C
ctrl-shRNA
SNX17-shRNA
50 KDa
37 KDa
SNX17
GAPDH
D
****
SNX17 levels (% ctrl-shRNA)
100
75
50
25
0
ctrl-shRNA
SNX17-shRNA
E
ctrl-shRNA
SNX17-shRNA
eGFP
SNX17
SNX17-R
75 KDa
37 KDa
GFP-SNX17
GAPDH
F
mEPSC Amplitude (pA)
30
20
10
0
**
****
ns
baseline
cLTP<30min
cLTP>30min
ctrl-shRNA
SNX17-shRNA
G
mEPSC Frequency (Hz)
12
10
8
6
4
2
0
*
**
ns
baseline
cLTP<30min
cLTP>30min
ctrl-shRNA
SNX17-shRNA

Figure S1. Validation of SNX17-shRNA, and effect of SNX17 knockdown in the initiation and maintenance of cLTP.

(A) Representative confocal images of DIV16 rat hippocampal neurons transfected at DIV12 with eGFP (filler) and either scrambled non-target shRNA SHC002 (Millipore Sigma) or SNX17-shRNA (TRCN0000190340, Millipore Sigma). Neurons were fixed, permeabilized and incubated with an anti-SNX17 antibody.

(B) Identification of two independent shRNA clones (clone1: TRCN0000190340 and clone 2: TRCN0000382281) to knockdown rat SNX17. The shRNA clones were used to generate lentiviruses and infect Rat2 cells. 5 days post infection, cell lysates were analyzed by western blot. pLKO.1 scrambled non-target shRNA SHC002 (Millipore Sigma) was used as a control for SNX17 levels. GAPDH was used as a loading control.

(C) Rat cortical neurons were infected with either SNX17-shRNA (clone 1) or control-shRNA at an MOI of 2. 6 days post-infection, cell extracts were collected and analyzed by western blot. GAPDH was used as a loading control.

(D) The levels of SNX17 protein were quantified in neurons infected with SNX17-shRNA (clone 1), and normalized to SNX17 levels in control-shRNA-infected neurons. ctrl-shRNA: 100%; SNX17-shRNA: 11.420 ± 0.852%. N=3 independent experiments. Statistical significance was determined using unpaired two-tailed Student's t-test, ****$p<0.001$. Error bars are SEM.

(E) HEK293 cells stably expressing the tet repressor (HEK293-TR) were either transfected with control-shRNA or SNX17-shRNA in the absence or presence of eGFP, GFP-SNX17 or shRNA-resistant GFP-SNX17 (SNX17-R), as indicated. 5-days post-infection, cells were treated with 1 μg/ml of doxycycline to promote the expression of eGFP, GFP-

SNX17 or GFP-SNX17-R. 24 hours later, extracts were collected and analyzed by western blot.

(F) DIV16-18 rat hippocampal neuron cultures transfected at DIV12 with a scramble control shRNA (ctrl-shRNA) or SNX17-shRNA, and either treated with cLTP or left untreated (baseline). mEPSCs were recorded during the first 30 min after cLTP (cLTP<30) or from 30 to 90 min after cLTP (cLTP>30), and mEPSC amplitude was quantified. ctrl-shRNA baseline: 11.890 ± 0.413, N=11 neurons; ctrl-shRNA cLTP<30min: 17.370 ± 1.655, N=9 neurons; ctrl-shRNA cLTP>30min: 15.730 ± 1.004, N=9 neurons; SNX17-shRNA baseline: 11.930 ± 0.488, N=21 neurons; SNX17-shRNA cLTP<30 min: 12.650 ± 0.555, N=13 neurons; SNX17-shRNA cLTP>30 min: 13.160 ± 0.497, N=12 neurons. 3 independent experiments. Data were analyzed by one-way ANOVA with Tukey's post hoc test, **p<0.01, ****p<0.001. Error bars are SEM.

(G) Same as F, but mEPSC frequency was quantified. ctrl-shRNA baseline: 0.888 ± 0.127, N=11 neurons; ctrl-shRNA cLTP<30min: 2.729 ± 0.844, N=9 neurons, ctrl-shRNA cLTP>30min: 2.051 ± 0.754, N=9 neurons, SNX17-shRNA baseline: 0.719 ± 0.116, N=21 neurons, SNX17-shRNA cLTP<30 min: 0.927 ± 0.114, N=13 neurons; SNX17-shRNA cLTP>30 min: 1.583 ± 0.327, N=12 neurons. 3 independent experiments. Data were analyzed by one-way ANOVA with Tukey's post hoc test, *p<0.05, ****p<0.001. Error bars are SEM.

**Figure S2**

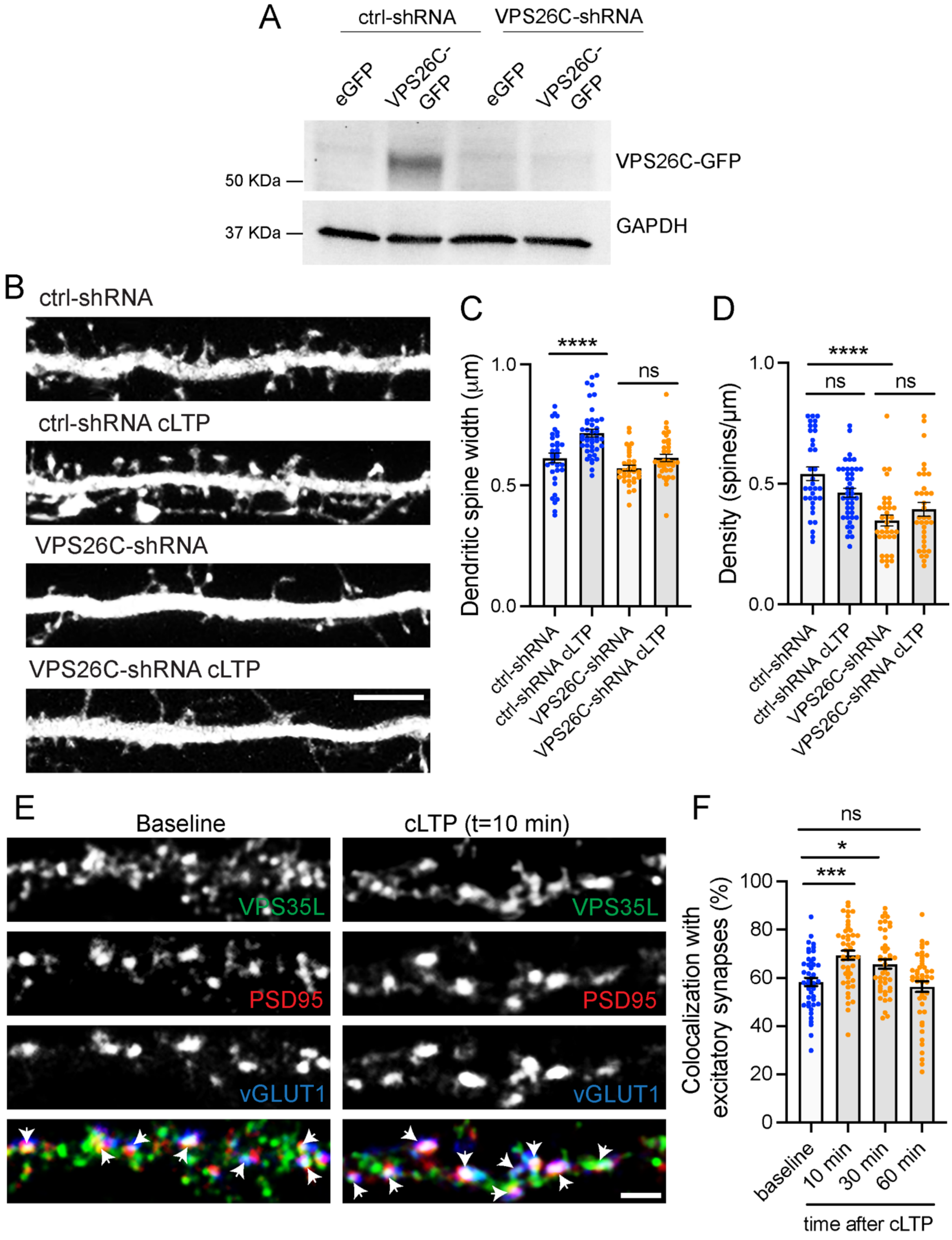
A
ctrl-shRNA
VPS26C-shRNA
eGFP
VPS26C-GFP
eGFP
VPS26C-GFP
VPS26C-GFP
GAPDH
50 KDa
37 KDa
B
ctrl-shRNA
ctrl-shRNA cLTP
VPS26C-shRNA
VPS26C-shRNA cLTP
C
Dendritic spine width (μm)
1.0
0.5
0.0
****
ns
ctrl-shRNA
ctrl-shRNA cLTP
VPS26C-shRNA
VPS26C-shRNA cLTP
D
Density (spines/μm)
1.0
0.5
0.0
****
ns
ns
ctrl-shRNA
ctrl-shRNA cLTP
VPS26C-shRNA
VPS26C-shRNA cLTP
E
Baseline
cLTP (t=10 min)
VPS35L
PSD95
vGLUT1
F
Colocalization with excitatory synapses (%)
100
80
60
40
20
0
ns
*
***
baseline
10 min
30 min
60 min
time after cLTP

Figure S2. The Retriever complex is necessary for the cLTP-dependent increase in dendritic spine width, and is dynamically recruited to dendritic spines upon cLTP.

(A) Validation of VPS26C-shRNA by western blotting. HEK293 cells stably expressing the tet repressor (HEK293-TR) were either transfected with control-shRNA (RHS4346, Horizon Discovery) or VPS26C-shRNA (V3LMM_455807, Horizon Discovery) in the absence or presence of eGFP or VPS26C-GFP, as indicated. 5-days post-infection, cells were treated with 1 μg/ml of doxycycline to promote the expression of eGFP or VPS26C-GFP. 24 hours later, extracts were collected and analyzed by western blot.

(B) Representative confocal images of dendritic spines in DIV16 hippocampal neurons co-transfected at DIV12 with eGFP (filler) and either ctrl-shRNA or VPS26C-shRNA. Neurons were either treated with cLTP or left untreated, and fixed 50 min after cLTP. Scale bar, 5 μm.

(C) The maximum width for each spine was quantified, and the average size of the dendritic spines in the first 50 μm of secondary dendrites was calculated. ctrl-shRNA: 0.656 ± 0.015, N=34 neurons; ctrl-shRNA with cLTP: 0.733 ± 0.019, N=42 neurons; VPS26C-shRNA: 0.581 ± 0.011, N=34 neurons; VPS26C-shRNA with cLTP: 0.626 ± 0.015, N=34 neurons. 3 independent experiments. Data were analyzed by one-way ANOVA with Tukey's post hoc test, ****$p<0.001$. Error bars are SEM.

(D) Quantification of spine density (spines/μm). ctrl-shRNA: 0.600 ± 0.034, N=34 neurons; ctrl-shRNA with cLTP: 0.458 ± 0.022, N=42 neurons; VPS26C-shRNA: 0.349 ± 0.025, N=34 neurons; VPS26C-shRNA with cLTP: 0.401 ± 0.030, N=34 neurons. 3 independent experiments. Data were analyzed by one-way ANOVA with Tukey's post hoc test, ****$p<0.001$. Error bars are SEM.

(E) Representative confocal images showing the colocalization of VPS35L with excitatory synapses before and 10 min after cLTP treatment. DIV19 rat hippocampal neuron cultures were untreated or treated with cLTP for 5 min ($Mg^{2+}$−free HBS with: 400 μM glycine, 20 μM bicuculline, and 3 μM strychnine). Cells were fixed after cLTP at 10, 30, or 60 min, permeabilized and incubated with antibodies against VPS35L, PSD95 and vGLUT1. Arrows indicate example of colocalization. Scale bar, 2 μm.

(F) The percentage of excitatory synapses that colocalize with VPS35L at the different time points was determined using Mander's colocalization coefficient (x100). Baseline: 58.330 ± 1.725, N=45; 10 min after cLTP: 69.380 ± 1.917, N=45; 30 min after cLTP: 65.730 ± 1.898, N=44; 60 min after cLTP: 56.380 ± 2.228, N=45. 3 independent experiments. Data were analyzed by one-way ANOVA with Tukey's post hoc test, $*p<0.05$, $***p<0.005$. Error bars are SEM.

**Figure S3**

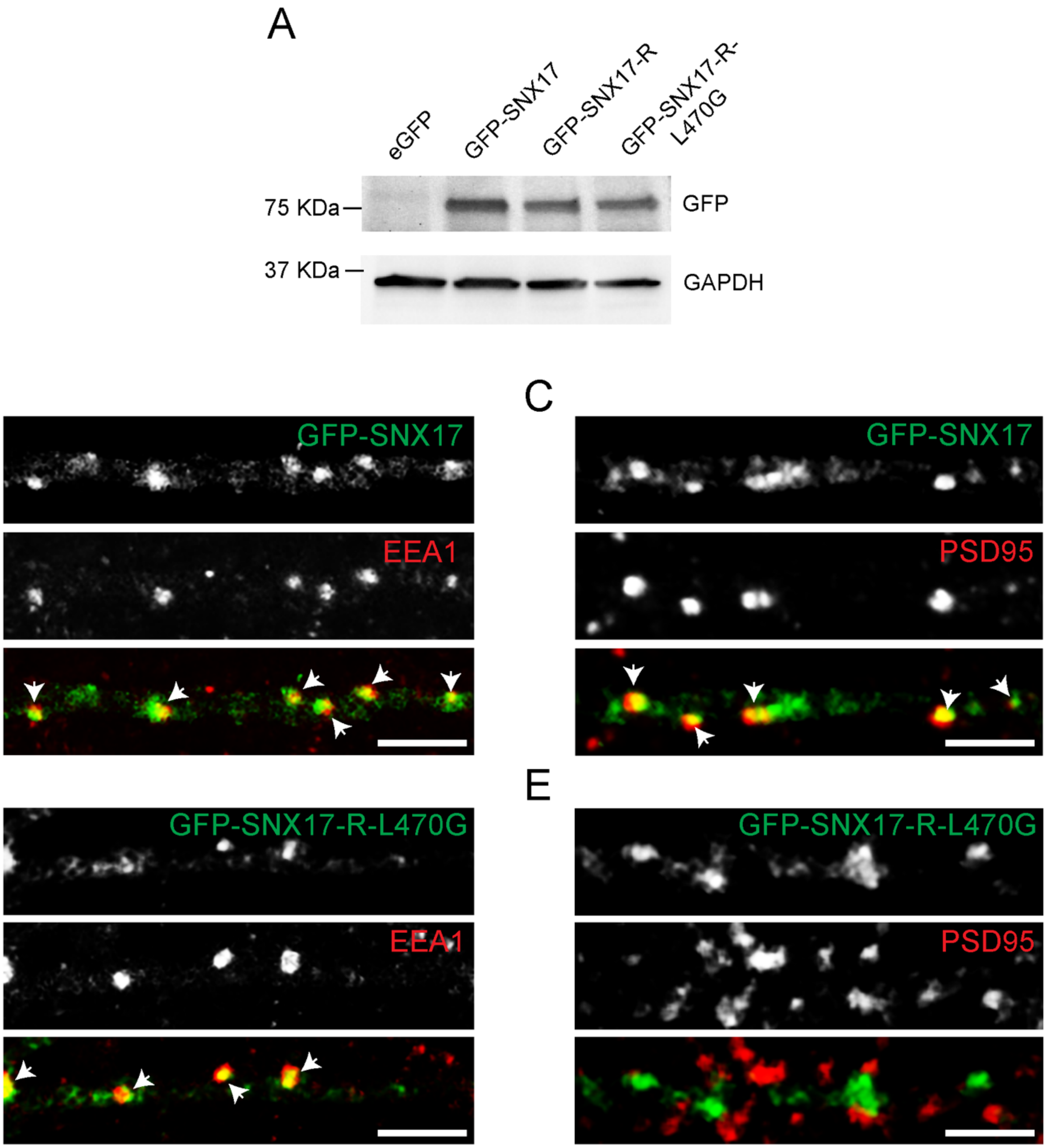
A
eGFP
GFP-SNX17
GFP-SNX17-R
GFP-SNX17-R-L470G
75 KDa
GFP
37 KDa
GAPDH
B
GFP-SNX17
EEA1
C
GFP-SNX17
PSD95
D
GFP-SNX17-R-L470G
EEA1
E
GFP-SNX17-R-L470G
PSD95

Figure S3. Validation of GFP-SNX17 and GFP-SNX17-L470G constructs

(A) HEK293T cells were transfected with eGFP, GFP-SNX17, GFP-SNX17-R or GFP-SNX17-R-L470G. 48 h later, extracts were collected and analyzed by western blot with an anti-GFP antibody. GAPDH was used as a loading control.

(B) Rat hippocampal neurons were transfected at DIV16 with GFP-SNX17, fixed 24 hours later, and stained for EEA1. EEA1 was labeled with Alexa Fluor-405, and pseudo colored in red to facilitate visualization. Arrows: examples of colocalization. Scale bar, 5 μm.

(C) Rat hippocampal neurons were transfected at DIV16 with GFP-SNX17, fixed 24 hours later, and stained for PSD95. PSD95 was labeled with Alexa Fluor-405, and pseudo colored in red to facilitate visualization. Arrows indicate examples of colocalization. Scale bar, 5 μm.

(D) Rat hippocampal neurons were transfected at DIV16 with GFP-SNX17-R-L470G, fixed 24 hours later, and stained for EEA1. EEA1 was labeled with Alexa Fluor-405, and pseudo colored in red to facilitate visualization. Arrows: examples of colocalization. Scale bar, 5 μm.

(E) Rat hippocampal neurons were transfected at DIV16 with GFP-SNX17, fixed 24 hours later, and stained for PSD95. PSD95 was labeled with Alexa Fluor-405, and pseudo colored in red to facilitate visualization. Arrows indicate examples of colocalization. Scale bar, 5 μm.

**Figure S4**

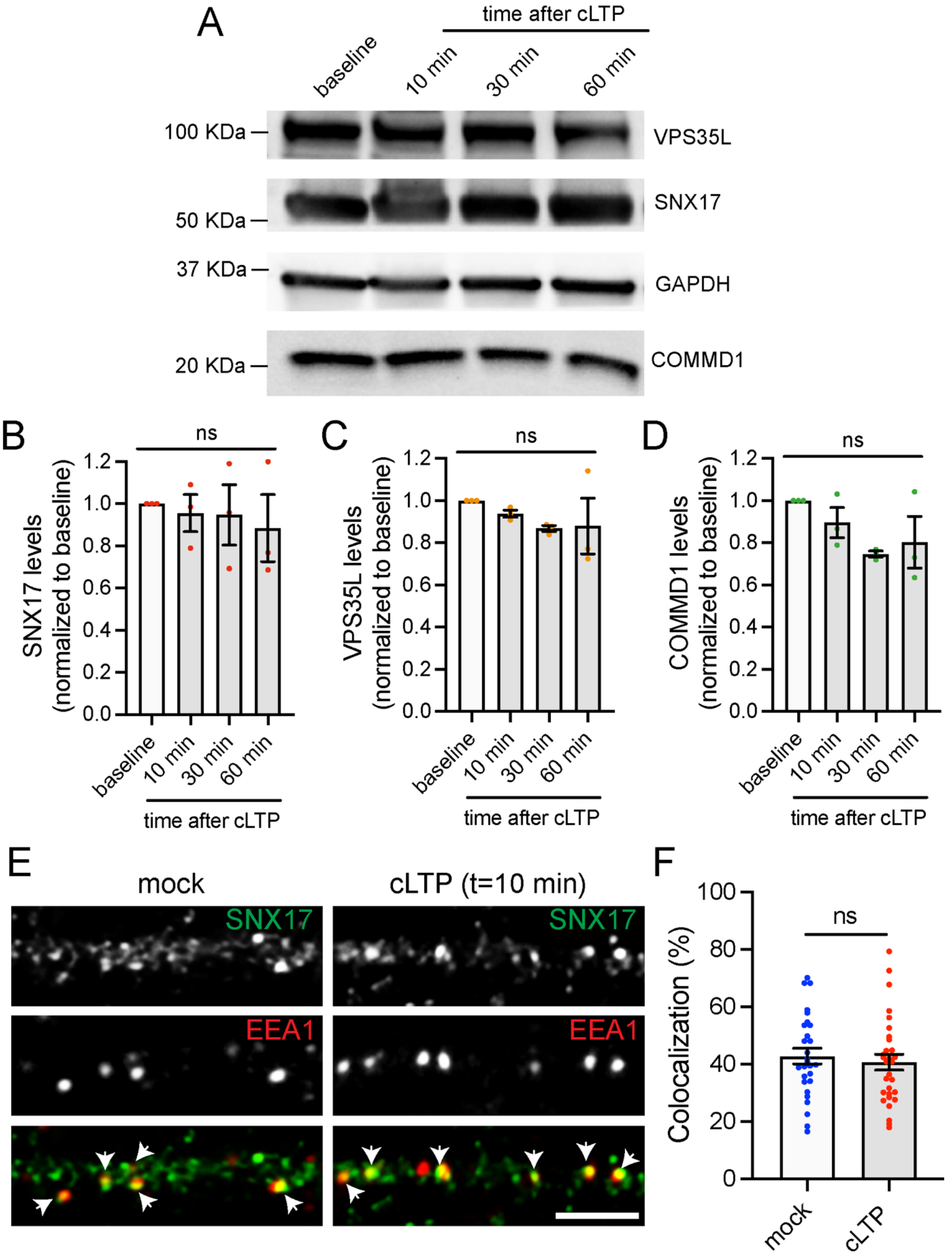
A
time after cLTP
baseline
10 min
30 min
60 min
100 KDa
VPS35L
SNX17
50 KDa
37 KDa
GAPDH
20 KDa
COMMD1
B
ns
SNX17 levels (normalized to baseline)
1.2
1.0
0.8
0.6
0.4
0.2
0.0
baseline
10 min
30 min
60 min
time after cLTP
C
ns
VPS35L levels (normalized to baseline)
D
ns
COMMD1 levels (normalized to baseline)
E
mock
cLTP (t=10 min)
SNX17
EEA1
F
ns
Colocalization (%)
100
80
60
40
20
0
mock
cLTP

Figure S4. Quantification of the total levels of proteins of the SNX17-Retriever pathway upon cLTP, and colocalization of SNX17 with EEA1 in the presence or absence of cLTP.

(A) DIV17 rat cortical neurons were either left untreated (baseline) or treated with cLTP for 5 min. Extracts were generated at 10, 30, or 60 min after cLTP, and analyzed by western blot for the levels of VPS35L, SNX17, COMMD1, and GAPDH as a loading control.

(B) The levels of SNX17 at 10, 30, or 60 min after cLTP were quantified and normalized to the baseline protein levels. Baseline: 1.000; 10 min: 0.955 ± 0.088; 30 min: 0.947 ± 0.144; 60 min: 0.885 ± 0.159. N=3 independent experiments. Data were analyzed by one-way ANOVA. Error bars are SEM.

(C) The levels of VPS35L at 10, 30, or 60 min after cLTP were quantified and normalized to the baseline protein levels. Baseline: 1.000; 10 min: 0.939 ± 0.017; 30 min: 0.868 ± 0.014; 60 min: 0.879 ± 0.132. N=3 independent experiments. Data were analyzed by one-way ANOVA. Error bars are SEM.

(D) The levels of COMMD1 at 10, 30, or 60 min after cLTP were quantified and normalized to the baseline protein levels. Baseline: 1.000; 10 min: 0.896 ± 0.071; 30 min: 0.747 ± 0.015; 60 min: 0.803 ± 0.123. N=3 independent experiments. Data were analyzed by one-way ANOVA. Error bars are SEM.

(E) DIV17 hippocampal neurons were treated with cLTP or HBS control (mock), washed, and incubated in HBS for 10 min following the cLTP (or mock) stimulus. Neurons were then fixed and stained for SNX17 and the early endosomal marker EEA1. Arrows indicate examples of colocalization. Scale bar, 5 µm.

(F) The percentage of EEA1 that colocalizes with SNX17 was quantified using Mander's colocalization coefficient (x100). Mock-treated neurons: 42.850 ± 2.723, N=28; cLTP-treated neurons: 40.730 ± 2.789, N=30. 3 independent experiments. Data were analyzed by unpaired two-tailed Student's t-test. Error bars are SEM.

**Figure S5**

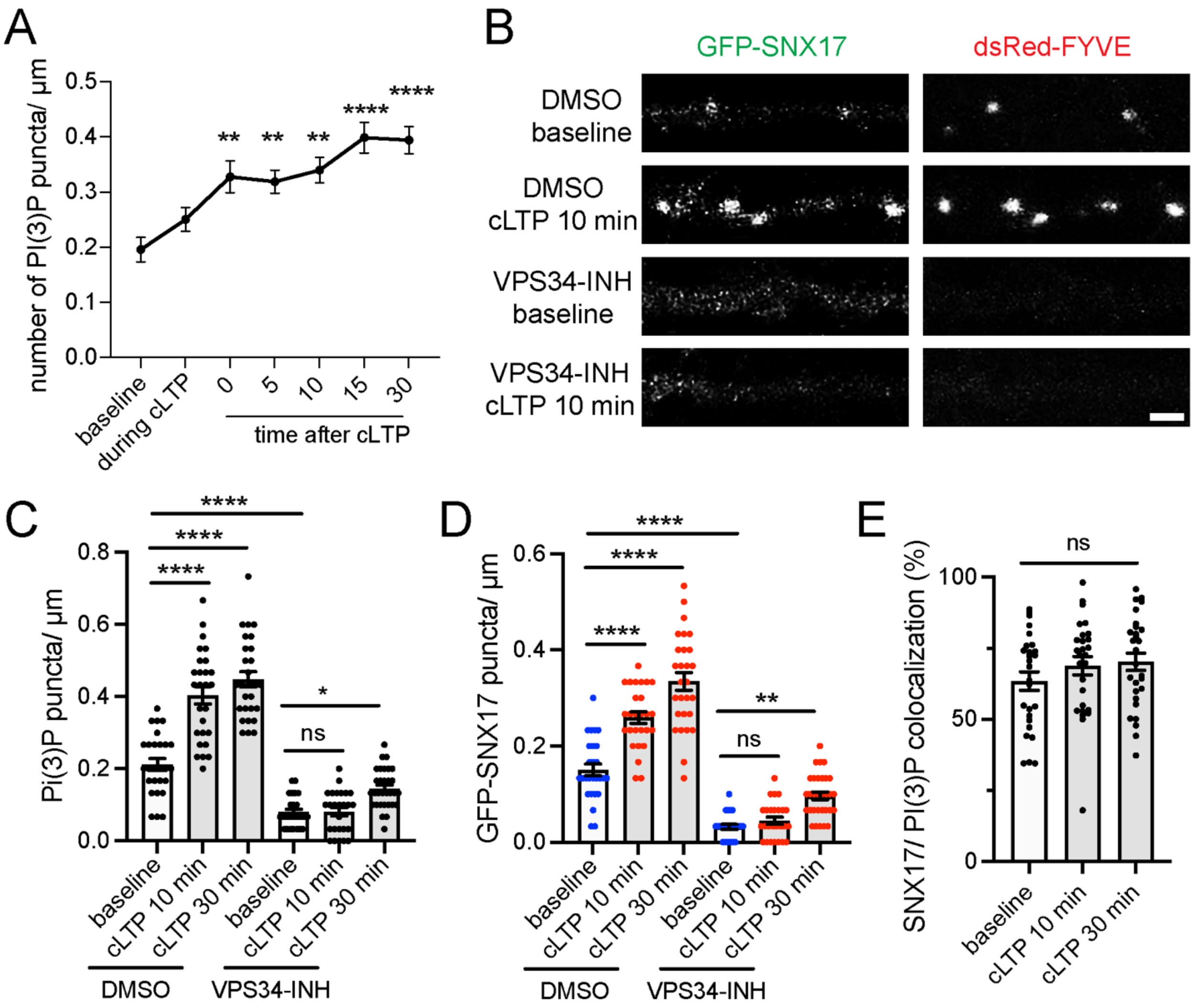

A
number of PI(3)P puncta/ µm
0.5
0.4
0.3
0.2
0.1
0.0
**
**
**
****
****
baseline
during cLTP
0
5
10
15
30
time after cLTP
B
GFP-SNX17
dsRed-FYVE
DMSO baseline
DMSO cLTP 10 min
VPS34-INH baseline
VPS34-INH cLTP 10 min
C
Pi(3)P puncta/ µm
0.8
0.6
0.4
0.2
0.0
****
****
****
ns
*
baseline
cLTP 10 min
cLTP 30 min
baseline
cLTP 10 min
cLTP 30 min
DMSO
VPS34-INH
D
GFP-SNX17 puncta/ µm
0.6
0.4
0.2
0.0
****
****
****
ns
**
baseline
cLTP 10 min
cLTP 30 min
baseline
cLTP 10 min
cLTP 30 min
DMSO
VPS34-INH
E
SNX17/ PI(3)P colocalization (%)
100
50
0
ns
baseline
cLTP 10 min
cLTP 30 min

Figure S5. Increased PI(3)P synthesis upon cLTP regulates SNX17 recruitment to dendritic spines as well as the formation of extrasynaptic SNX17-positive puncta.

(A) Rat hippocampal neurons were transfected at DIV16 with dsRed-FYVE. 24 h later, neurons were either left untreated (baseline) or treated with cLTP for 5 min and fixed at the indicated time points after the cLTP stimulus. Baseline: 0.196 ± 0.022, N=31 neurons; during cLTP: 0.250 ± 0.021, N=30 neurons; 0 min after cLTP: 0.328 ± 0.029, N=30 neurons; 5 min after cLTP: 0.319 ± 0.021, N=30 neurons; 10 min after cLTP: 0.340 ± 0.023, N=30 neurons; 15 min after cLTP: 0.399 ± 0.028, N=30 neurons; 30 min after cLTP: 0.394 ± 0.025, N=30 neurons. 3 independent experiments. Data were analyzed by one-way ANOVA with Tukey's post hoc test, **p<0.01, ****p<0.001. Error bars are SEM.

(B) Representative confocal images of DIV17 hippocampal neurons that were transfected at DIV16 with GFP-SNX17 and dsRed-FYVE. 24 h later, neurons were treated with either DMSO or 1 μM VPS34-INH for 15 min. Neurons were either fixed in the absence of cLTP (baseline) or treated with cLTP for 5 min, and fixed 10 or 30 min after the cLTP stimulus. DMSO or VPS34-INH were maintained during the course of the experiment. Images are shown for the baseline and 10 min after cLTP time points in the presence or absence of VPS34-INH. Scale bar, 2 μm.

(C) The number of dsRed-FYVE- positive puncta in the first 30 μm of secondary dendrites was quantified at the indicated time points. Baseline DMSO: 0.212 ± 0.016, N=27 neurons; cLTP 10 min DMSO: 0.404 ± 0.024, N=27 neurons; cLTP 30 min DMSO: 0.448 ± 0.021, N=28 neurons; baseline VPS34-INH: 0.079 ± 0.008, N=26 neurons; cLTP

10 min VPS34-INH: 0.081 ± 0.011, N=26 neurons; cLTP 30 min VPS34-INH: 0.145 ± 0.010, N=31 neurons. 3 independent experiments. Statistical significance was determined using one-way ANOVA with Tukey's post hoc test, *p<0.05, ****p<0.001. Error bars are SEM.

(D) The number of GFP-SNX17 puncta in the first 30 μm of secondary dendrites was quantified at the indicated time points. Baseline DMSO: 0.151 ± 0.012, N=27 neurons; cLTP 10 min DMSO: 0.259 ± 0.012, N=27 neurons; cLTP 30 min DMSO: 0.335 ± 0.019, N=28 neurons; baseline VPS34-INH: 0.032 ± 0.005, N=26 neurons; cLTP 10 min VPS34-INH: 0.045 ± 0.007, N=26 neurons; cLTP 30 min VPS34-INH: 0.097 ± 0.008, N=31 neurons. 3 independent experiments. Statistical significance was determined using one-way ANOVA with Tukey's post hoc test, **p<0.01, ****p<0.001. Error bars are SEM.

(E) The colocalization between dsRed-FYVE- and GFP-SNX17- positive puncta, in the absence of VPS34-INH, was analyzed using Mander's colocalization coefficient (x100). Baseline: 63.430 ± 3.248%, N=27 neurons; cLTP 10 min: 68.850 ± 3.258%, N=27 neurons, cLTP 30 min: 70.260 ± 3.060%, N=28 neurons. Data were analyzed using one-way ANOVA with Tukey's post hoc test. Error bars are SEM.
